# Sexually dimorphic alternative splicing and its erosion following the loss of sex in stick insects

**DOI:** 10.64898/2026.02.21.707240

**Authors:** Iulia Darolti, Marjorie Labédan, Vincent Mérel, Darren J. Parker, Tanja Schwander

## Abstract

Sexually dimorphic traits emerge from divergent selective pressures acting on males and females and are facilitated by gene regulatory mechanisms. While gene expression differences have been the subject of considerable focus in the evolution of sexual dimorphism, other regulatory processes remain comparatively understudied. One such mechanism is alternative splicing, whereby a single gene can encode multiple distinct transcripts, leading to functional diversification of the proteome. Leveraging long-read PacBio Iso-sequencing, we examine patterns of sex differences in gene expression and alternative splicing in three tissues of related *Timema* stick insect species, including five sexual and four asexual lineages. Alternative splicing is widespread in *Timema*, with the greatest isoform complexity found in the male gonad tissue. We observe similar levels of sex differences in gene expression and splicing, and a core set of genes involved in sexual differentiation and germline development that have conserved differential splicing patterns across the *Timema* phylogeny. However, patterns of sex-biased alternative splicing undergo faster lineage-specific turnover than sex-biased gene expression, and differentially spliced genes are subject to strong pleiotropic and purifying constraints. Following the transition from sexual reproduction to asexuality, there is a systematic reduction in isoform diversity and erosion of sex-biased splicing variation. Taken together, our findings suggest a role for sexual selection in maintaining splicing complexity in sexual *Timema* species.

## Introduction

A central aim in evolutionary biology is to understand how a single genome can generate multiple phenotypes, such as queen and worker castes in eusocial insects (Price et al., 2018), morphologically and ecologically distinct larval, pupal and adult stages of holometabolous insects (Simpson et al., 2011), or different sexes, which represent one of the most extreme and prevalent forms of intra-specific diversity (Mank, 2017). Owing to their distinct reproductive strategies, males and females are often subject to opposing selective pressures, which can generate conflicts in a shared genome (Lande, 1980). Intralocus conflicts occur when traits with a shared genetic basis have opposing fitness effects in different selective contexts, as is the case for sexually antagonistic traits in males and females (Bonduriansky & Chenoweth, 2009). These conflicts can potentially be resolved through regulatory evolution, ultimately leading to the formation and maintenance of sexually dimorphic traits (Rowe et al., 2018).

Differential gene expression is a major mechanism by which selection mitigates the constraints of a shared genome (Connallon & Knowles, 2005; Mank, 2017) and a large body of research has investigated the role of sex-biased genes in sex-specific adaptation (Darolti & Mank, 2023; Djordjevic et al., 2022; Harrison et al., 2015; Immonen et al., 2017; Ingleby et al., 2014; Magnusson et al., 2011; Perry et al., 2014; Whittle & Extavour, 2019). Such studies have greatly advanced our understanding of sexually dimorphic traits by identifying candidate loci and signaling pathways (Emlen et al., 2012; Galouzis & Prud’homme, 2021; Kopp et al., 2000; Toubiana et al., 2021). However, gene expression levels represent only one dimension of phenotypic diversification, and the contribution of other regulatory mechanisms to facilitating the evolution of distinct male and female forms has been comparatively understudied.

One such regulatory mechanism is post-transcriptional alternative splicing which can generate structurally variable isoforms of a gene, for example through different combinations of exon exclusion or intron retention events (Verta & Jacobs, 2022). This process expands transcriptome and proteome complexity (Nilsen & Graveley, 2010; Tseng et al., 2022) and is known to regulate important biological functions such as tissue development (Wang et al., 2008), diverse physiological processes (Kalsotra & Cooper, 2011; Tseng et al., 2022), and sex-determination (Schutt & Nothiger, 2000). However, a significant proportion of isoforms are likely non-functional, largely representing stochastic mis-splicing (Benitiere et al., 2024; Sherin et al., 2026; Wan & Larson, 2018). Alternative splicing rates can vary substantially across phenotypes (Jacobs & Elmer, 2021; Price et al., 2018), tissues (Naftaly et al., 2021), developmental stages (Gibilisco et al., 2016), and species (Blekhman et al., 2010). As such, the extent to which this splicing diversity reflects functional innovation and phenotypic optimization as opposed to relaxed selection and noise remains an open question.

Evidence from diverse taxa indicates that sex-biased alternative splicing is widespread and extends well beyond primary sex-determination pathways (Blekhman et al., 2010; Chen et al., 2023; Gibilisco et al., 2016; McIntyre et al., 2006; Naftaly et al., 2021; Rogers et al., 2021; Singh & Agrawal, 2023). However, we still lack a phylogenetically explicit understanding of how sex-biased splicing evolves across lineages, how stable such patterns are through evolutionary time, and to what extent they are actively maintained by sexual selection rather than arising from neutral or stochastic processes. A powerful way to disentangle these forces is to compare closely related species that differ in reproductive mode and, consequently, in the presence of sexual selection and efficiency of purifying selection. The stick insect genus *Timema* provides such a natural evolutionary experiment, whereby multiple independent transitions from sexual reproduction with separate sexes to asexuality have occurred within the clade (Schwander et al., 2011). Asexual *Timema* species are morphologically and ecologically equivalent to their sexual sister species (Law & Crespi, 2002), however, because they consist of only females, they experience no sexual selection or sexual conflict (Parker et al., 2019b). Moreover, many of the former sexually dimorphic traits are expected to decrease in strength or disappear as they lose their adaptive value (Schwander et al., 2013). Interestingly, as *Timema* species possess an XX:X0 sex determination system, rare males can be produced in asexual lineages both through the accidental loss of an X chromosome during oogenesis (Schwander et al., 2013), and via rare sexual reproduction events. Such asexual males provide a unique control when investigating patterns of sex-biased transcriptomes, as the transcriptional profiles underlying male phenotypes in these lineages are expected to evolve primarily under relaxed selection and genetic drift.

Here, we leverage PacBio long-read Iso-sequencing (Iso-seq) data from somatic and reproductive tissues of males and females from five sexual and four asexual *Timema* lineages to provide an isoform-resolved analysis of sex-biased gene regulation. Specifically, we compare sex differences in gene expression and alternative splicing across tissues, species, and reproductive modes to test whether sex-biased splicing is conserved or labile, whether differentially spliced genes are subject to distinct selective constraints compared to sex-biased genes, and how these regulatory patterns change following the loss of sex and sexual selection. By integrating long-read transcriptomics with a replicated evolutionary framework, our study directly addresses the evolutionary forces shaping alternative splicing in the context of sexual dimorphism and sexual conflict.

## Materials and Methods

### Biological samples, Iso-seq library preparation and sequencing

We conducted PacBio Iso-seq of RNA previously extracted for different projects (Parker et al., 2026). We used RNA extractions of three tissues (femur, gut, and gonad) of males and females collected in the field as juveniles and reared to adulthood in the laboratory under standard conditions (see Djordjevic et al., 2025 for details). Prior to dissection, all individuals were fed with an artificial diet for two days, allowing their gut contents to be cleared of plant material (their natural food source). Insects were then anesthetized with CO_2_, dissected, and tissues individually flash-frozen in liquid nitrogen before being stored at −80°C. Extractions were done using a TRIzol-Chloroform protocol (Djordjevic et al., 2025). RNA quantity and quality was measured using a fluorescent RNA-binding dye (QuantiFluor RNA System) and a Fragment Analyzer (Agilent Technologies). Equimolar amounts of RNA extractions from two to four individuals were then pooled to generate a single Iso-seq library per tissue, sex, and species (for a total of 48 libraries, see Table S1). SMRTbell library preparation and Sequel II sequencing using two SMRT cells per sample were completed at the Lausanne Genomic Technologies Facility (University of Lausanne).

### Genome assemblies and annotations

Previously published genome assemblies for the five sexual *Timema* species (NCBI BioSamples SAMN41934017, SAMN48771607, SAMN48771608, SAMN48771609, SAMN48771610) (Djordjevic et al., 2025; Parker et al., 2026) were repolished with POLCA v4.1.4 (Zimin & Salzberg, 2020) and then annotated. Briefly, *Arthropoda* protein sequences from OrthoDB v11 (Kuznetsov et al., 2023) and species-specific RNA-seq data were used to annotate each genome assembly using BRAKER 3 v3.0.8 (Gabriel et al., 2024). UTRs were added with stringtie2utr.py from the BRAKER suite. The pipelines for genome assembly and annotation can be found at https://github.com/vpymerel/TimemaSex_AssemblyAnnotation.

### Iso-seq read processing, isoform identification and filtering

SMRT Cell subread data was run through the PacBio Iso-Seq Analysis pipeline in SMRT Link v.12, which generated full-length non-concatemer (FLNC) reads trimmed of primers and poly(A) tails. Through the same pipeline, FLNCs were then clustered to produce high quality consensus (HQ) transcripts, with 99% sequence accuracy and supported by a minimum of two FLNCs.

HQ transcript sequences from all sexual and asexual samples in a species pair (Table S1, Table S2) were aligned to the genome assembly and annotation of the sexual species using pbmm2 v1.10 (https://github.com/PacificBiosciences/pbmm2), with the options --preset ISOSEQ, --sort, --unmapped. Redundant transcripts were collapsed into unique isoforms using isoseq3 collapse v3.8.2. To avoid a high rate of reference transcript redundancy, we set --max-5p-diff 1000000 and --max-3p-diff 1000000 and collapsed isoforms that matched a reference transcript at all splice junctions but had an alternative 5’ or 3’ end, which may reflect partial cDNA synthesis or degradation (Tardaguila et al., 2018).

Isoform characterization and filtering was performed using SQANTI3 v5.1.1 (Pardo-Palacios et al., 2024). For each species, we used previously generated male and female RNA-seq short-read data from the same three tissues (femur, gut, gonad) (Parker et al., 2026) to provide information on isoform expression and splice junction coverage. Specifically, we aligned short-reads to the non-redundant isoform sequences with kallisto v0.48 (Bray et al., 2016) to obtain expression matrices and separately aligned short-reads to the reference genome and non-redundant isoform annotation with STAR v2.7.11a (Dobin et al., 2013) to obtain splice junction coordinates and coverage. Based on this information, we used SQANTI3 to remove isoforms classified as intra-priming products, RT-switching artifacts, and isoforms with non-canonical junctions that have short-read coverage < 3. We further excluded mono-exonic transcripts and collapsed isoforms with the same open reading frame, only preserving the longest transcript for subsequent analyses.

### Differential gene expression and differential transcript usage analysis

For each species and tissue, we first mapped the paired-end short-read RNA sequencing data to the final filtered set of isoforms using Salmon v1.10.2 (Patro et al., 2017) with 100 bootstraps and correcting for GC and sequence-specific biases with --seqBias and --gcBias. We then extracted gene-level and transcript-level abundance matrices using the tximport v1.34 package (Soneson et al., 2015) in R v4.4.3 (R Core Team). Finally, for each tissue of each species we removed lowly expressed genes and isoforms by imposing a minimum 2 TPM and, respectively, 5 TPM expression threshold in at least half of the individuals of one sex.

We identified differentially expressed (DE) genes with the DESeq2 v1.46 package (Love et al., 2014) in R, with an adjusted false discovery rate < 0.05 and an absolute log_2_ fold change ≥ 1. To identify differentially spliced (DS) genes we used a differential transcript usage analysis. Unlike standard differential expression analysis, differential transcript usage tests whether the composition of transcripts changes across conditions by measuring shifts in within-gene relative proportion of each isoform (the fraction of a gene’s total expression that is accounted for by each isoform), irrespective of whether total gene expression differs between conditions. For this, we first transformed transcript abundance estimates from Salmon to normalize for transcript length and library size using the dtuScaledTPM method in tximport. We then modeled within-gene relative isoform proportions using the Dirichlet-multinomial model from the DRIMSeq v1.34 R package (Nowicka & Robinson, 2016) with parameters min_samps_feature_expr=2, min_samps_feature_prop=2, min_samps_gene_expr=2, min_feature_expr=10, min_feature_prop=0.1, min_gene_expr=10. To reduce false positives and control the overall false discovery rate (Love et al., 2018), the resulting gene-and transcript-level *p* values were subjected to stage-wise adjustment with the stageR v1.28 package (Van den Berge et al., 2017). A first screening stage was performed at the gene level, with genes failing the adjusted *p* value threshold of 0.05 excluded from further analysis. Secondly, for genes passing the first stage, a transcript-level significance testing was performed.

### Gene orthology inference

For each species, we used the nucleotide sequences for the filtered set of isoforms and selected the longest isoform per-gene. Orthologs between species were inferred using OrthoFinder v2.5.5 (Emms & Kelly, 2019) with default parameters.

### Estimating gene expression tissue specificity

Tissue specificity of gene expression is best estimated across many different tissues. Therefore, in addition to the short-read RNA-seq data for the three focal tissues with long-read sequencing data (femur, gonad, and gut), we also included short-read data from antenna, brain, defense glands, and fat body from Parker et al. (2026), and estimated gene expression as detailed above. For each gene, we calculated a tissue specificity index (*τ*) (Yanai et al., 2005) based on the formula: 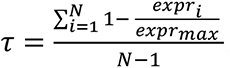, where *N* is the number of tissues, *expr* is the expression of the gene in tissue *i*, and *expr_max_* is the maximum expression of that gene across all tissues. *τ* values range from 0 to 1, where higher values indicate a higher tissue-specific expression, while lower values indicate a more even expression across tissues.

### Estimating rates of coding-sequence evolution

For all single-copy orthologs identified through the gene orthology inference, we extracted nucleotide coding-sequences as inferred by SQANTI3. We aligned sequences using macse v2.07 (Ranwez et al., 2018) and removed alignments shorter than 150bp after gap removal. We used the branch model test (model = 1, nssites = 0) in the CODEML package in PAML v4.8 (Yang, 2007) to estimate rates of coding-sequence evolution for each branch of the phylogeny “((*T. poppense*,*T. californicum*),*T. cristinae*),(*T. podura*,*T. bartmani*));“. To avoid divergence estimates being affected by mutational saturation and to exclude cases of gene misalignment or poor alignment, we removed genes with the rate of synonymous substitutions (*d*_S_) > 2 as per previous studies (Axelsson et al., 2008). For each category of genes (DE and DS genes), we extracted from the PAML results the number of synonymous (*D*_S_) and non-synonymous (*D*_N_) substitutions, as well as the number of synonymous (S) and non-synonymous (N) sites. For each category, we then calculated the average rate of synonymous substitutions (*d*_S_) and the average rate of non-synonymous substitutions (*d*_N_) as *d*_S_ = *D*_S_ / S and, respectively, *d*_N_ = *D*_N_ / N. Bootstrapping with 1,000 replicates was used to calculate the 95% confidence intervals for divergence estimates in each group. Lastly, to test for significant differences in *d*_N,_ *d*_S_, and *d*_N_/*d*_S_ between gene categories, we used a random subsampling test. Specifically, when comparing evolutionary rates between two gene sets, from the larger gene set we generated 1,000 random subsets without replacement, each to the size of the smaller gene set. For every random gene subset, we calculated *d*_N_ and *d*_S_ as described above, and *d*_N_/*d*_S_ as the ratio between the two. The observed evolutionary rate estimates for the smaller gene set were compared to the distribution of statistics obtained from the random subsets and two-sided adjusted *p* values were obtained.

## Results and Discussion

### Alternative splicing is prevalent in *Timema* stick insects

We discovered high levels of splicing variation in *Timema* stick insects, recovering between 18,686 and 30,212 isoforms across the nine focal comparisons (Table S2). These comprise four sexual-asexual species pairs (*T. poppense* + *T. douglasi*, *T. californicum* + *T. shepardi*, *T. cristinae* + *T. monikensis*, *T. podura* + *T. genevievae*), analyzed together on the sexual species reference, and one sexual species (*T. bartmani*) analyzed alone. Using a combination of PacBio Iso-seq and Illumina paired-end RNA-sequencing we found that 40-53% of genes in the curated *Timema* transcriptomes have multiple isoforms (Fig. S1). Our estimates of splicing variation are likely conservative as we imposed stringent filters to limit false positive isoforms. The proportion of alternatively spliced genes that we observe across *Timema* species is most directly comparable to those from recent long-read invertebrate studies, such as those in planthoppers (58%) (Tong et al., 2022) and honey bees (68%) (Zheng et al., 2023), which similarly highlight extensive transcriptomic complexity. In contrast, earlier studies using less sensitive short-read sequencing approaches reported generally lower rates, including in silkworms (27%) (Li et al., 2012), pea aphids (34%) (Grantham & Brisson, 2018), *Drosophila* (37%) (Gibilisco et al., 2016), and flatworms (42%) (Wang et al., 2015). This distinction highlights that reported fractions of alternatively spliced genes are often as much a reflection of sequencing sensitivity as they are of underlying biology. This pattern is also reflected in our data, where the species with the fewest samples (*T. bartmani*) also shows the lowest number of filtered isoforms (Table S1, S2).

### Transcriptional complexity varies across biological levels

Across all nine analyzed species, we observed similar patterns of spliceosome complexity variation among tissues and between sexes. At the tissue level, we found that the gonad has a significantly higher number of isoforms per-gene than both the femur (Wilcoxon rank-sum test, W = 9, *p* = 0.007) and the gut (Wilcoxon rank-sum test, W = 0, *p* < 0.001) (Fig. 1A, Fig. S2). Of the genes expressed in all three tissues, 50% have more than one isoform in the gonad, in contrast to 34% in the femur and the gut. A high splicing complexity in the gonad emerges as a systematic pattern across multiple clades (Gibilisco et al., 2016; Mazin et al., 2021; Naftaly et al., 2021), with brain and liver being the few somatic tissues with similarly high levels of splicing (Gibilisco et al., 2016; Grosso et al., 2008; Mazin et al., 2021). Compared to somatic tissues, gonads express the largest number of splicing regulators and exhibit strong tissue-specific expression of splicing-factor genes (Grosso et al., 2008). Together with the extreme heterogeneity in germ and somatic cells present in the gonad, this may underly the observed isoform diversity in the reproductive tissue.

**Figure 1.**
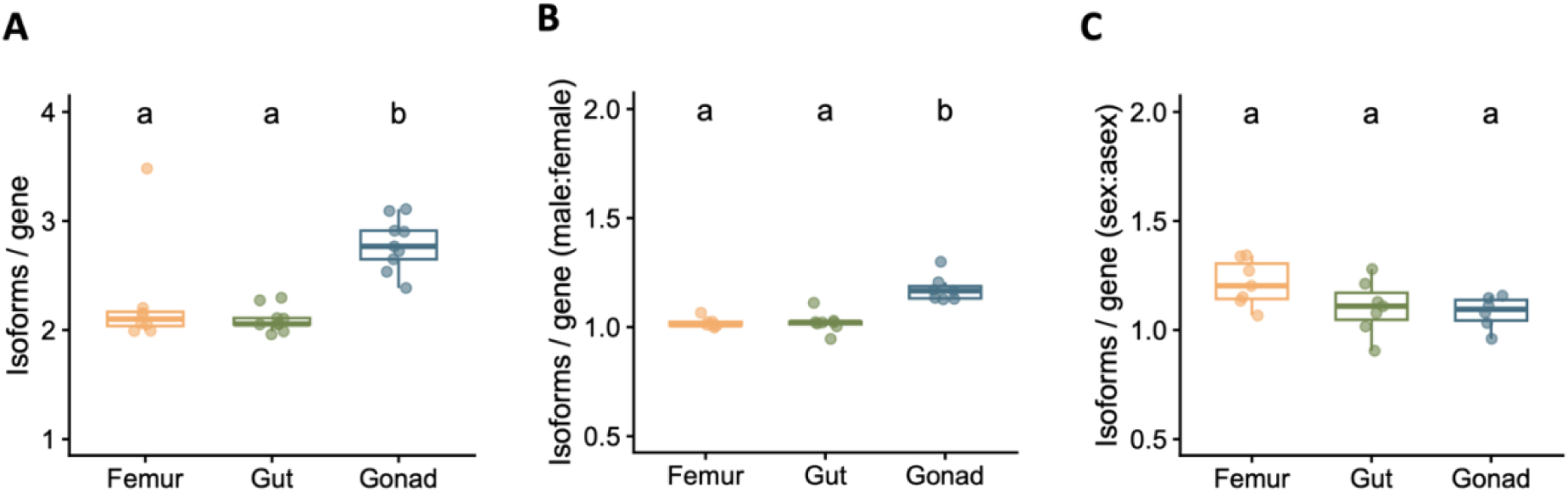
Transcriptome complexity across tissues, between sexes, and between reproductive modes. (A) Average number of isoforms per-gene for each femur, gut, and gonad sample of each species. Points represent species-specific values, and for each species, values were calculated from genes expressed in all three tissues and with more than one isoform in at least one of the tissues. (B) The ratio between males and females within each species and tissue in the average number of isoforms per-gene. Points represent species-specific values, and for each species, included are genes expressed in both sexes and with more than one isoform in at least one of the sexes. C) The ratio between sexual and asexual individuals in the average number of isoforms per-gene. Data points comprise sex-matched comparisons for each sexual-asexual pairs (sexual vs. asexual females, and sexual vs. asexual males). Included are genes expressed in both sexual and asexual individuals and with more than one isoform in at least one of the reproductive modes. In all panels, results from pairwise Wilcoxon rank-sum tests are represented by lowercase letters, where boxplots labelled with distinct letters are significantly different (*p*_adj_ < 0.05), while boxplots sharing a letter are non-significant.

Gonad spliceosome complexity further differs markedly between the sexes (Fig. 1B). In the reproductive tissues, the male-to-female ratio significantly exceeds parity for both the number of expressed genes (one-sample t-test against null = 1, *t*(6) = 11.5, *p*_adj_ < 0.001, 95% CI [1.11,1.17]; Fig. S3) and per-gene isoform diversity (*t*(6) = 7.6, *p*_adj_ < 0.001, 95% CI [1.12,1.23]; Fig. 1B, Fig. S4). These patterns of male bias are restricted to the gonads (Wilcoxon rank-sum tests, *p*_adj_ < 0.001 in all comparisons), as male:female ratios did not deviate from the null in either somatic tissue (*p*_adj_ > 0.05 in all comparisons).

This pattern contrasts with some previous reports in *Drosophila* species which suggested that males and females adopt different strategies for diversifying their transcriptome, with testes expressing more genes while a higher fraction of alternatively spliced genes being found in the ovary (Gibilisco et al., 2016). The male-bias in both isoform and gene expression diversity that we observe here may reflect the pronounced cellular heterogeneity of the testis, which recent single-cell atlases have repeatedly identified as one of the most transcriptionally and cell-type diverse tissues (Han et al., 2018; Han et al., 2020; Li et al., 2022). Alternatively, these results may arise from a more permissive chromatin environment of spermatocytes and spermatids during meiosis and spermiogenesis (Soumillon et al., 2013). Such an open chromatin state could also promote stochastic splicing variation. This could occur by increasing transcriptional permissiveness or by reducing the fidelity of splice-site recognition under conditions of elevated transcriptional output and saturation of the splicing machinery (Mazin et al., 2021). Although the higher isoform diversity in *Timema* male gonads is not accompanied by a disproportionate increase in non-coding relative to coding isoforms (Fig. S5), the male-biased pattern of isoform diversity is notably chromosome-dependent. Gonad expressed X-linked genes show in fact reduced isoform diversity in males relative to autosomal genes (Fig. S6), a finding that is consistent with meiotic X chromosome inactivation in the male germline of *Timema* species (Djordjevic et al., 2025). Future coupling of alternative splicing data with single-cell transcriptomics or chromatin accessibility analysis could help disentangle between these factors. Alternatively, sperm competition has been implicated in driving rapid testis transcriptome evolution (Trost et al., 2023) and, to some extent, this might also accelerate splicing rates in males (Rogers et al., 2021), although comparative tests at the isoform diversity level are lacking.

Although alternative splicing is widespread across eukaryotes, the extent to which such events are functional and adaptive, as opposed to reflecting splicing noise, remains unclear (Wan & Larson, 2018). Isoform diversity tends to scale with organismal complexity (Chen et al., 2014), consistent with a role of alternative splicing in expanding functional regulatory repertoires. At the same time, this pattern does not exclude a role for non-adaptive processes, as under the “drift barrier” theory, reduced effective population size (*Ne*) can increase the accumulation of deleterious mutations, including mutations that impair splicing fidelity, thereby increasing isoform diversity through splicing errors (Benitiere et al., 2024). Indeed, an analysis across dozens of metazoan species revealed an inverse correlation between rates of alternative splicing and proxies for *Ne*, a pattern driven predominantly by low-abundance and non-functional splicing variants (Benitiere et al., 2024).

Our analyses focused on high-confidence isoforms retained after filtering for likely technical and biological artifacts. Consequently, low-abundance variants and potential mis-splicing events were largely excluded. Among the retained high-confidence isoforms, however, our results do not support a simple relationship between population size and isoform diversity. Autosomal *Ne* varies by more than tenfold across sexual *Timema* species, ranging from ∼162,000 in *T. poppense* to ∼1,900,000 in *T. podura* (Parker et al., 2022), yet we detected no significant correlation between *Ne* estimates and alternative splicing levels (Fig. S7). To control for differences in sequencing depth among species, we randomly subsampled 400,000 FLNCs from each library and then repeated the complete isoform identification and filtering analysis based on these subsampled datasets. Additionally, to ensure that estimates of alternative splicing levels are not confounded by differences in the number of expressed genes, we restricted the analysis to the gonad and gut tissues, since femur was missing from *T. bartmani*, and to genes expressed in all five sexual species. The absence of a correlation between *Ne* and isoform diversity cannot therefore be attributable to differences in the number of input reads, nor the number of expressed genes. Additionally, lowly expressed genes are expected to exhibit greater isoform diversity, as low expression may increase the frequency of stochastic or erroneous splicing events (Benitiere et al., 2024). However, within our set of isoforms, we did not recover a negative relationship between gene expression and the number of isoforms per gene (Fig. S8), further suggesting that the isoform diversity retained in our analyses is not predominantly driven by low-abundance, stochastic splicing noise.

The lack of effective recombination in asexual *Timema* species is expected to significantly reduce the efficacy of selection, leading to an accelerated accumulation of deleterious mutations (Bast et al., 2018). This relaxed constraint can decrease the efficiency of the spliceosome and increase mis-splicing events (Benitiere et al., 2024) in asexual compared to sexual species. Contrary to this expectation, we find a significantly greater isoform diversity in sexual *Timema* species compared to asexual species in both femur (one-sample t-test against null = 1, *t*(6) = 5.4, *p*_adj_ = 0.005, 95% CI [1.11,1.31]) and gonad (*t*(5) = 2.7, *p*_adj_ = 0.04, 95% CI [1.00,1.16]), but not in gut (*t*(6) = 2.2, *p*_adj_ = 0.07, 95% CI [0.99,1.22]) (Fig. 1C, Fig. S9).

The magnitude of the sexual:asexual ratio, however, did not differ significantly among tissues (Wilcoxon rank-sum tests, *p*_adj_ > 0.05 in all comparisons). Taken together, these results indicate that, among high-confidence isoforms, splicing variation in sexual *Timema* is not primarily explained by genetic drift or stochastic mis-splicing, but likely reflects a regulated and potentially functional layer of transcriptomic complexity.

### Conservation and turnover of sex differences in splicing

Alternative splicing differences between the sexes have commonly been regarded as secondary to sex-biased gene expression. Our analyses of global patterns of differential gene expression and differential transcript usage between males and females across the five *Timema* sexual species suggest instead a comparable prevalence of these two regulatory mechanisms. Consistently across species, we find the gonad tissue to be highly transcriptionally dimorphic at both the gene expression level and the splicing level, while somatic tissues show only minimal differentiation (Fig. 2). Although we identify twice as many differentially expressed (DE) genes as differentially spliced (DS) genes in the gonad, these rates converge when accounting for the total number of filtered genes that could be included in each analysis, with, on average, 32% sex-biased genes and 34% genes with differential splicing. In stark contrast, sexual dimorphism in the femur and gut was minor, with only 1-2% DE genes and 4-7% DS genes. Our finding that gonads exhibit profound sexual dimorphism across both regulatory levels is consistent with previous research (Blekhman et al., 2010; Gibilisco et al., 2016; Naftaly et al., 2021; Rogers et al., 2021; Singh & Agrawal, 2023). However, while sex-biased splicing has been viewed as a more minor component of transcriptional dimorphism compared to differential gene expression (Gibilisco et al., 2016; Rogers et al., 2021), the fact that we find a comparable prevalence between these two mechanisms suggests that alternative splicing can also be a fundamental driver of sex-specific phenotypic differentiation. Our detection of such a high proportion of genes with differential transcript usage may stem from the application of long-read sequencing, which captures full-length isoforms that are often undetected with short-read RNA-sequencing pipelines. The functional contribution of these two regulatory mechanisms to sex-specific differentiation warrants further validation.

**Figure 2.**
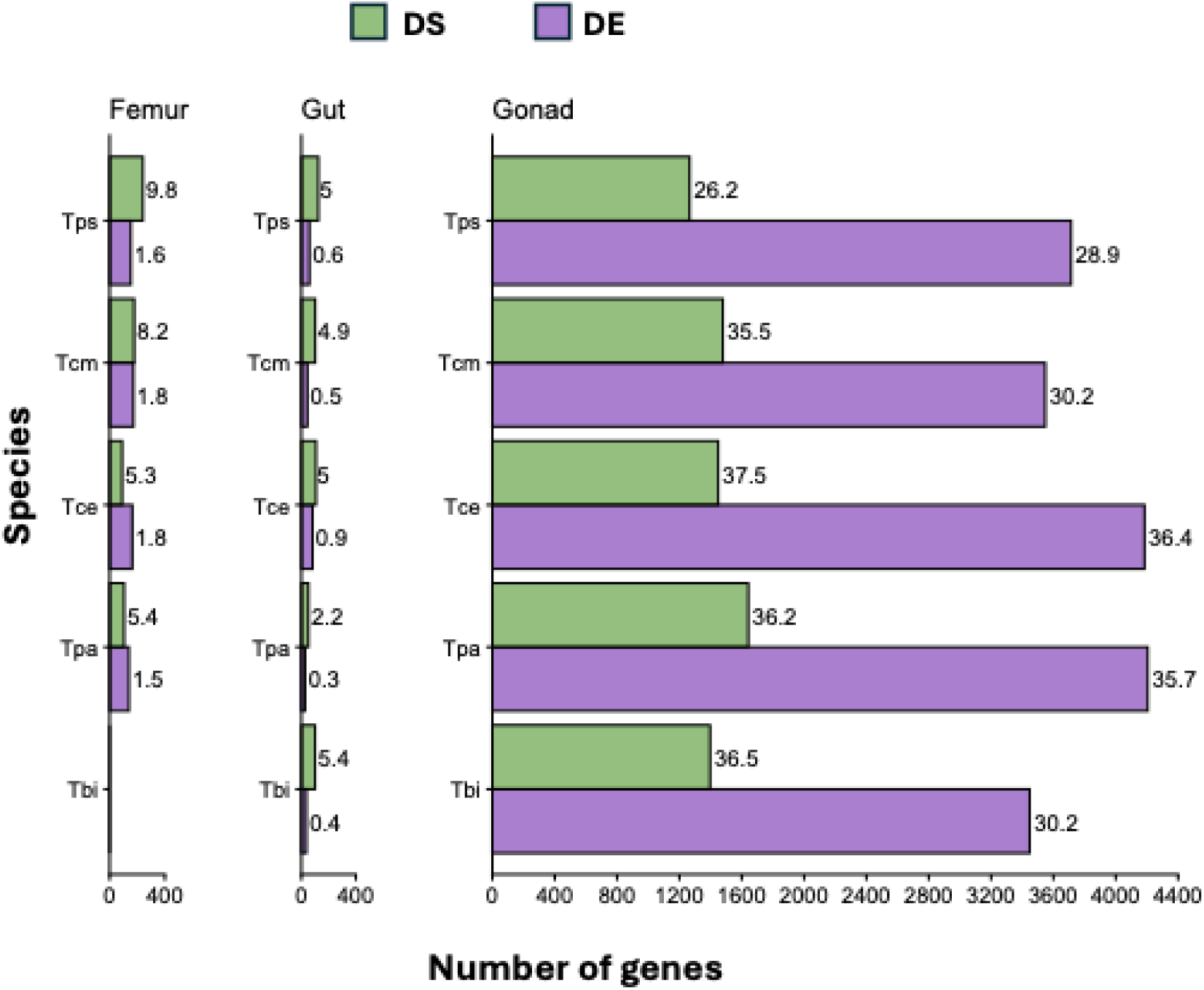
Differential gene expression and differential transcript usage between males and females in *Timema* sexual species. Number of genes with a significant difference in expression (DE) and in transcript usage patterns (DS) between males and females in femur, gut, and gonad tissues of *T. poppense* (Tps), *T. californicum* (Tcm), *T. cristinae* (Tce), *T. podura* (Tpa) and *T. bartmani* (Tbi). The numbers next to the bars represent the percentage of DE and DS genes out of the total number of genes that could be tested in each respective analysis.

Sex-biased gene expression can exhibit rapid evolutionary turnover across species and lineages (Grath & Parsch, 2016; Harrison et al., 2015). In contrast, the conservation versus turnover of sex-biased alternative splicing remains comparatively unexplored. Here, we present the first comparative assessment of these dynamics across multiple species within a well-defined phylogeny. The proportion of DS genes are remarkably consistent across the five *Timema* sexual species (Fig. 2), which prompted us to investigate the extent to which these patterns of gonadal differential regulation are conserved across the phylogeny or represent independent lineage-specific events. We identified 6,311 one-to-one orthologous genes across the five species. Based on this orthologous gene set, we find significant all-species conservation in both DE genes (SuperExactTest *p*_adj_ < 0.001, 54-fold enrichment over random expectation) and DS genes (SuperExactTest *p*_adj_ < 0.001, 5-fold enrichment) (Fig. 3A, B). Part of this overlap may reflect inheritance of ancestral regulatory states, rather than independent maintenance by selection. Nonetheless, among the 54 genes with a conserved differential splicing pattern in all five species, we find several major splicing regulators and genes involved in germline development (Table 1). These results indicate that at least a core subset of sex-biased splicing events in sexual *Timema* species are functionally driven and involved in sexual differentiation, rather than a by-product of stochastic noise or neutral retention of ancestral splicing states.

**Figure 3.**
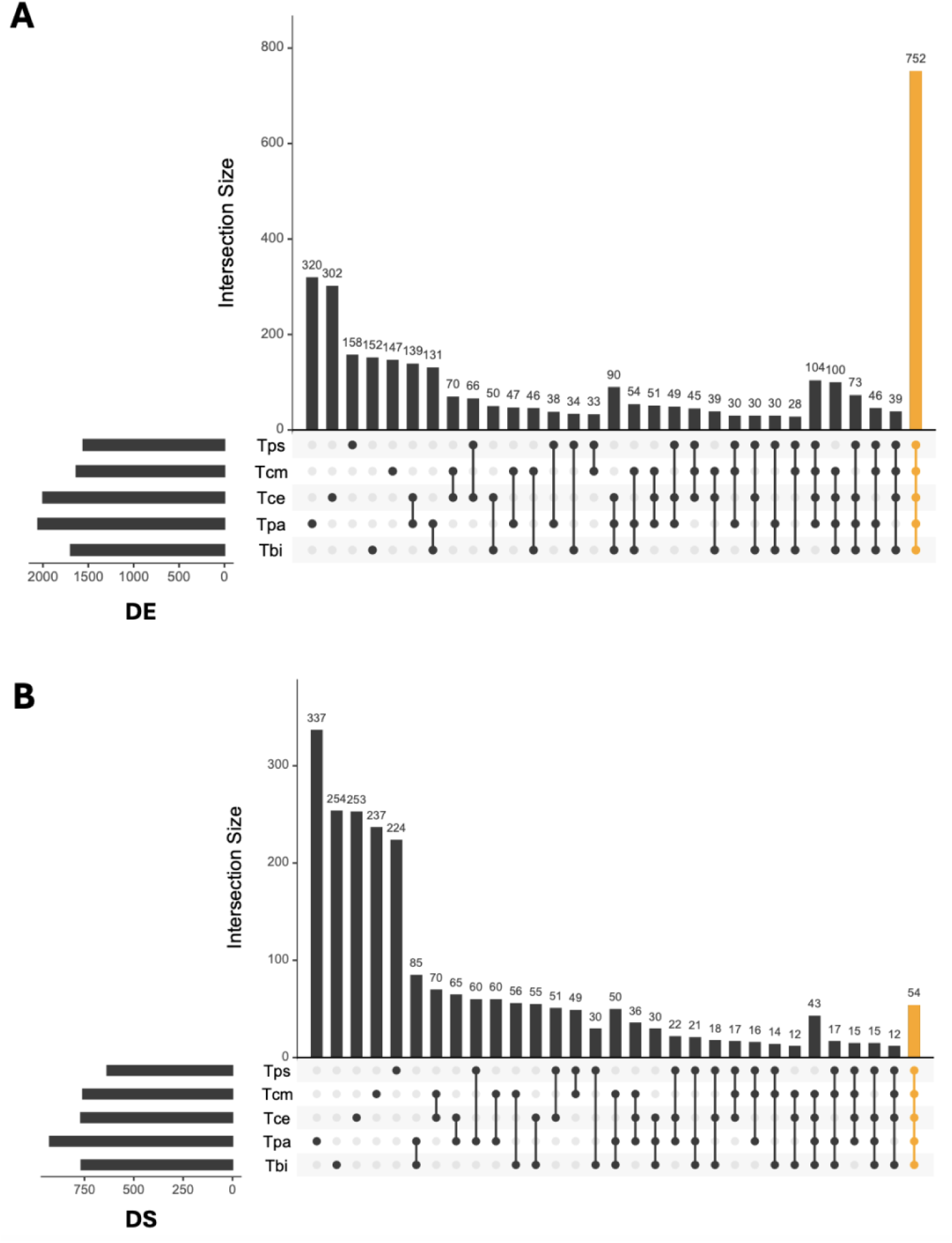
Intersection of orthologous gonad genes that are (A) differentially expressed (DE genes) and (B) genes with different transcript usage (DS genes) between males and females, across the five sexual species. The number of DE and DS genes shared across all species is shown in orange. Species name abbreviations: Tps = *T. poppense*, Tcm = *T. californicum*, Tce = *T. cristinae*, Tpa.= *T. podura*, Tbi = *T. bartmani*.

**Table 1.** Selected genes with conserved differential splicing pattern across the *Timema* phylogeny. Homology was established using BLASTp against the NCBI Arthropoda and *Drosophila* nr databases.

| Top Hexapoda blast hit | Top *Drosophila* blast hit | Characterized function in *Drosophila* |
| --- | --- | --- |
| homeotic protein female sterile isoform (*Periplaneta americana*, E-value 0.0, 100% query coverage, 65% identity) | fs(1)h, homeotic protein female sterile isoform (E-value 4e-89, 53% query coverage, 60% identity) | splicing coactivator involved in somatic sex determination (Qiu et al., 2019) |
| serine/arginine-rich splicing factor 6 (*Cryptotermes secundus* E-value 1e-115, 62% query coverage, 90% identity) | B52 (E-value 4e-87, 62% query coverage, 69% identity) | splicing factor; expressed in ovaries and developing eggs (Fear et al., 2015; Kraus & Lis, 1994) |
| muscleblind-like protein 3 (*Periplaneta americana*, E-value 0.0, 43% query coverage, 90% identity) | MBL, muscleblind (E-value 7e-170, 96% query coverage, 76% identity) | major regulator of alternative splicing; involved in tissue differentiation (Bargiela et al., 2014) |
| putative helicase MOV-10 (*Bacillus rossius redtenbacheri*, E-value 0.0, 98% query coverage, 54% identity) | ARMI, Armitage (E-value 1e-101, 64% query coverage, 35% identity) | involved in the localization of Piwi into Yb bodies (Saito et al., 2010) |
| anoctamin-10 (*Bacillus rossius redtenbacheri*, E-value 0.0, 95% query coverage, 76% identity) | AXS, abnormal X segregation (E-value 2e-118, 85% query coverage, 36% identity) | chromosome segregation during meiosis (Whyte et al., 1993) |
| protein ECT2-like isoform (*Macrosteles quadrilineatus*, E-value 0.0, 100% query coverage, 51% identity) | PBL, pebble (E-value 0.0, 81% query coverage, 52% identity) | role in cytokinesis (Deduyer et al., 2025) |
| poly(1) RNA polymerase gld-2 homolog A (*Anabrus simplex*, E-value 0.0, 100% query coverage, 59% identity) | GLD2 (E-value 4e-129, 54% query coverage, 56% identity) | mRNA translation control; regulator of late spermatogenesis (Sartain et al., 2011) |

While genes with higher isoform counts were more likely to show conserved differential splicing and those with fewer isoforms tended to have a species-specific differential splicing pattern (Fig S10A), this difference in transcript complexity does not account for the observed cross-species conservation. Even when restricting the analysis to genes with at most three isoforms, DS genes still show a significant overlap among species (Fig. S10B; SuperExactTest *p*_adj_ < 0.001, 10-fold enrichment). Thus, the conservation of DS genes is unlikely to be driven simply by high transcript complexity at a small number of loci, and instead appears to reflect a stable regulatory program.

Sex-biased genes, and in particular male-biased genes, frequently exhibit accelerated evolutionary rates at both the coding-sequence and expression levels (Grath & Parsch, 2016). This rapid divergence has been attributed to intense sexual selection acting in males (Proschel et al., 2006; Sawyer et al., 2007), however, relaxed pleiotropic constraints may also facilitate these transitions (Dapper & Wade, 2020; Djordjevic et al., 2022; Harrison et al., 2015). Consistent with these patterns, we observe that while a significant core of both male-biased and female-biased genes are shared across *Timema* species (SuperExactTest *p*_adj_ < 0.001; Fig. S11), a substantial fraction of the sex-biased transcriptome is still species-specific or subject to rapid change. While high rates of expression turnover have been documented in diverse taxa, including *Drosophila* (Whittle & Extavour, 2019), Galliform birds (Harrison et al., 2015), mosquitoes (Papa et al., 2017), and fish (Lichilin et al., 2021), the evolutionary stability of sex-biased splicing has remained largely unexplored. Our results reveal a striking disparity in conservation between these two regulatory mechanisms, where the enrichment for conserved sex-biased gene expression (54-fold enrichment) is ten times greater than that for differential splicing (5-fold enrichment). This confirms that alternative splicing undergoes faster lineage-specific turnover (Barbosa-Morais et al., 2012; Gibilisco et al., 2016; Merkin et al., 2012; Rogers et al., 2021).

Alternative splicing may therefore be more evolutionarily labile than gene-level expression, enabling faster lineage-specific diversification of proteomes. This higher turnover could reflect either a greater flexibility of splicing regulation or a larger mutational target size. Differential gene expression is predominantly shaped by promoters, enhancers, and other distal regulatory elements, whereas alternative splicing may rely on a more dispersed cis-regulatory architecture, spanning splice sites, exonic and intronic enhancers and silencers across the gene body (Wang & Burge, 2008; Yeo et al., 2007). Together, these features may render sex-biased splicing more prone to lineage-specific regulatory divergence and therefore less evolutionarily conserved than sex-biased gene expression.

### Differentially expressed and spliced genes are under distinct constraints

An emerging pattern from genome-wide analyses suggests that transcriptional and splicing regulatory mechanisms are largely non-redundant as they target different gene sets and affect distinct biological processes (Grantham & Brisson, 2018; Jacobs & Elmer, 2021; Rogers et al., 2021; Singh & Agrawal, 2023; Smout et al., 2026). In the context of sexual dimorphism, it has been proposed that sex-biased gene expression and sex-biased alternative splicing may resolve sexual conflict through distinct molecular strategies (Rogers et al., 2021; Singh & Agrawal, 2023). Specifically, while sex-biased expression shifts the overall gene expression level towards a sex-specific optimum, sex-biased splicing can produce functional transcript variants with different fitness effects in the two sexes from a shared genomic locus. Therefore, alternative splicing may provide a powerful mechanism to resolve sexual conflict, particularly for genes under pleiotropic constraints (Rogers et al., 2021).

Throughout the *Timema* phylogeny, although we find that the overlap between DE and DS genes in the gonad tissue is not significantly different than expected by chance (hypergeometric tests, *p* > 0.05), on average 68% of DS genes have an unbiased expression between males and females (Fig. 4A). Moreover, among the 54 genes with a conserved sex-biased splicing pattern, only six genes also show a pattern of sex-biased gene expression in all the species. Our results are consistent with previous findings that, to a large extent, these two regulatory mechanisms impact different sets of genes.

**Figure 4.**
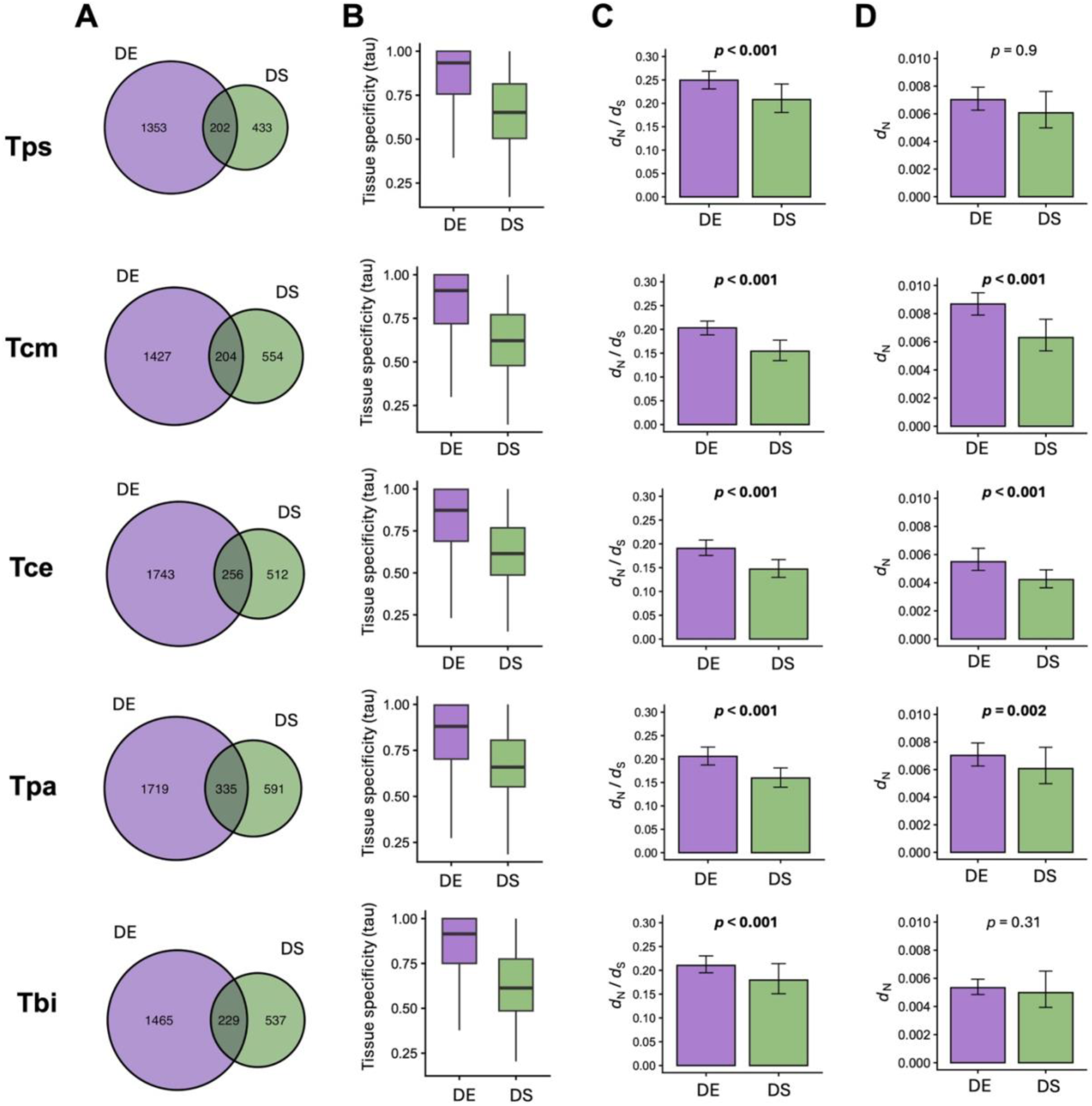
Overlap, tissue specificity, and rates of coding-sequence evolution for DE and DS genes in the gonad tissue of sexual *Timema* species. (A) Venn diagrams showing overlap between DE and DS genes. (B) Tissue specificity (tau) estimates based on expression information from seven tissues, where values closer to 1 represent higher tissue-specific expression, while values closer to 0 represent broader expression across tissues. In all comparisons, DS genes have a significantly lower tau value compared to DE genes as estimated by Wilcoxon rank-sum tests. (C) Estimates of the ratio between nonsynonymous (*d*_N_) and synonymous substitutions (*d*_S_). (D) Estimates of nonsynonymous substitution rates. In both (C) and (D) 95% confidence intervals are shown based on bootstrapping with 1,000 replicates, and adjusted *p* values are based on random subsampling tests with 1,000 replicates. Species name abbreviations: Tps = *T. poppense*, Tcm = *T. californicum*, Tce = *T. cristinae*, Tpa.= *T. podura*, Tbi = *T. bartmani*.

We additionally employed two measures for testing whether DE and DS genes are subject to different selective constraints. First, we used expression data across a range of tissues to obtain tissue specificity estimates. The evolution of sex-biased gene expression can be limited by pleiotropic constraints, as genes expressed across multiple tissues or developmental stages are often unbiased in expression and subject to stabilizing selection to maintain their functions in both males and females (Mank et al., 2008; Meisel, 2011). Alternative splicing may circumvent these pleiotropic constraints on gene expression level by generating distinct male and female isoforms (Rogers et al., 2021). In line with this, we observe that DS genes are significantly more broadly expressed than DE genes (Fig. 4B; Wilcoxon rank-sum tests, *p*_adj_ < 0.05 in all comparisons). Secondly, we calculated rates of coding-sequence evolution and found that DS genes have a lower *d*_N_/*d*_S_ than DE genes, attributable to a significantly lower rate of non-synonymous substitutions (Fig. 4C, D). To test whether the lower *d*_N_/*d*_S_ of DS genes was confounded by expression level, we compared *d*_N_/*d*_S_ across gene expression quartiles (Fig. S12). A two-way ANOVA test confirmed that gene category remained highly significant (*F*_1,1916_ = 35.92, *p* = 2.45 x 10^-9^) with no significant main effect of expression quartile (*p* = 0.24) or interaction between factors (*p* = 0.72). Within-quartile Wilcoxon rank-sum tests showed that DS genes exhibit significantly lower *d*_N_/*d*_S_ across moderate and high expression tiers (Q2-Q4: all *p*_adj_ < 0.01), with a non-significant trend only in the lowly expressed tier where DS genes were scarce (Q1: *p*_adj_ = 0.32). Therefore, the lower *d*_N_/*d*_S_ for DS genes is robust and independent of expression level. Taken together, our results suggest that, compared to differential expression, differential splicing more commonly occurs in genes under stronger purifying selection and functional constraints.

### Erosion of sex-biased splicing patterns in asexual species

The comparison between sexual and asexual lineages provides a natural experiment to disentangle selection, drift, and developmental constraint. The transition from sexual reproduction to asexuality significantly alters the selective landscape, which can lead to genome-wide transcriptomic changes (Bast et al., 2018; Liu et al., 2014; Parker et al., 2019a). Although reduced selective efficacy in asexual lineages might predict random drift of gene expression away from its optima (Huylmans et al., 2021), empirical evidence reveals that more complex selective dynamics are at work. Asexual lineages are relieved from sexual selection and sexual conflict, and this resolution should in theory allow asexuals to invest more towards female-specific traits and functions, potentially manifesting as large-scale transcriptomic shifts, such as the upregulation of female-biased genes or downregulation of formerly male-biased ones (Parker et al., 2019b). Contrary to this expectation, studies including those in *Timema* have found repeated patterns of desexualization of gene expression in asexual females, characterized by a lowered expression for female-biased genes, consistent with a change in female trait optima in asexual lineages and the decay of sexual traits (Huylmans et al., 2021; Parker et al., 2019b). Comparative studies of sexual and asexual lineages can thus provide a powerful test for determining whether patterns of sex differences in splicing are maintained by sexual selection. Moreover, rare asexual males, some which can arise sporadically through loss of an X chromosome in X0 systems, provide a unique opportunity to examine the consequences of transition to asexuality for male-associated splicing patterns as male-specific phenotypes are not expected to experience sustained selection in predominantly female lineages.

For the two sexual – asexual species pairs (*T. poppense* – *T. douglasi*; *T. podura* – *T. genevievae*) for which we have gonadal isoform information for both sexes, we find that, on average, 47% of DS genes between sexual males and females are also differentially spliced in the asexual lineages, an overlap which is significantly greater than expected by chance (fold enrichment = 1.2, *p* < 0.01, one-tailed hypergeometric test; background genes N = 4,129 for *T. poppense* – *T. douglasi* and N = 3,902 for *T. podura* – *T. genevievae*). These sex differences in splicing could represent broad developmental constraints that persist even after the transition from sexual to asexual reproduction. Nevertheless, a large fraction of DS genes found in the sexual lineages do not exhibit sex-biased transcript usage in the asexual lineages. As such, to further understand how patterns of sex-biased splicing change following the transition to asexuality, we conceptualized three evolutionary scenarios for how within-gene isoform proportions in asexual females (*F*_asex_) could relate to sex bias in the ancestral sexual lineage (Δ [*F*_asex_ – *M*_sex_]; Fig. 5A). Asexual females may largely show a conserved pattern of ancestral splicing regulation under stabilizing selection or developmental constrain, where sexual female-biased isoforms (red, Δ > 0) maintain a high per-gene isoform proportion (*F*_asex_ > 0.5), while male-biased isoforms (blue, Δ < 0) remain at low proportions (*F*_asex_ < 0.5). Conservation of splicing patters would also be expected if insufficient time has passed since the sexual and asexual lineages split to allow changes in splicing to accumulate. Relaxed selection or drift may lead to an erosion of sexual female patterns, wherein female-biased isoforms shift towards lower within-gene proportions in asexual females (Δ > 0, *F*_asex_ ≤ 0.5). Alternatively, and consistent with trends found at the gene expression level (Parker et al., 2019b), there may be a reversal in differential splicing patterns towards the opposite sex profile, with an increased proportion of male-biased isoforms in asexual females (Δ < 0, *F*_asex_ ≥ 0.5). Such a reversal could occur if male-biased isoforms provide some functional advantage in the new asexual context. Inverse patterns are expected for asexual males, whereby pattern conservation would predict retention of high male-biased isoform usage, pattern erosion would predict a loss of male-specific splicing patterns, and pattern reversal would predict an increase in within-gene relative proportion for female-biased isoforms (Fig. S13A).

**Figure 5.**
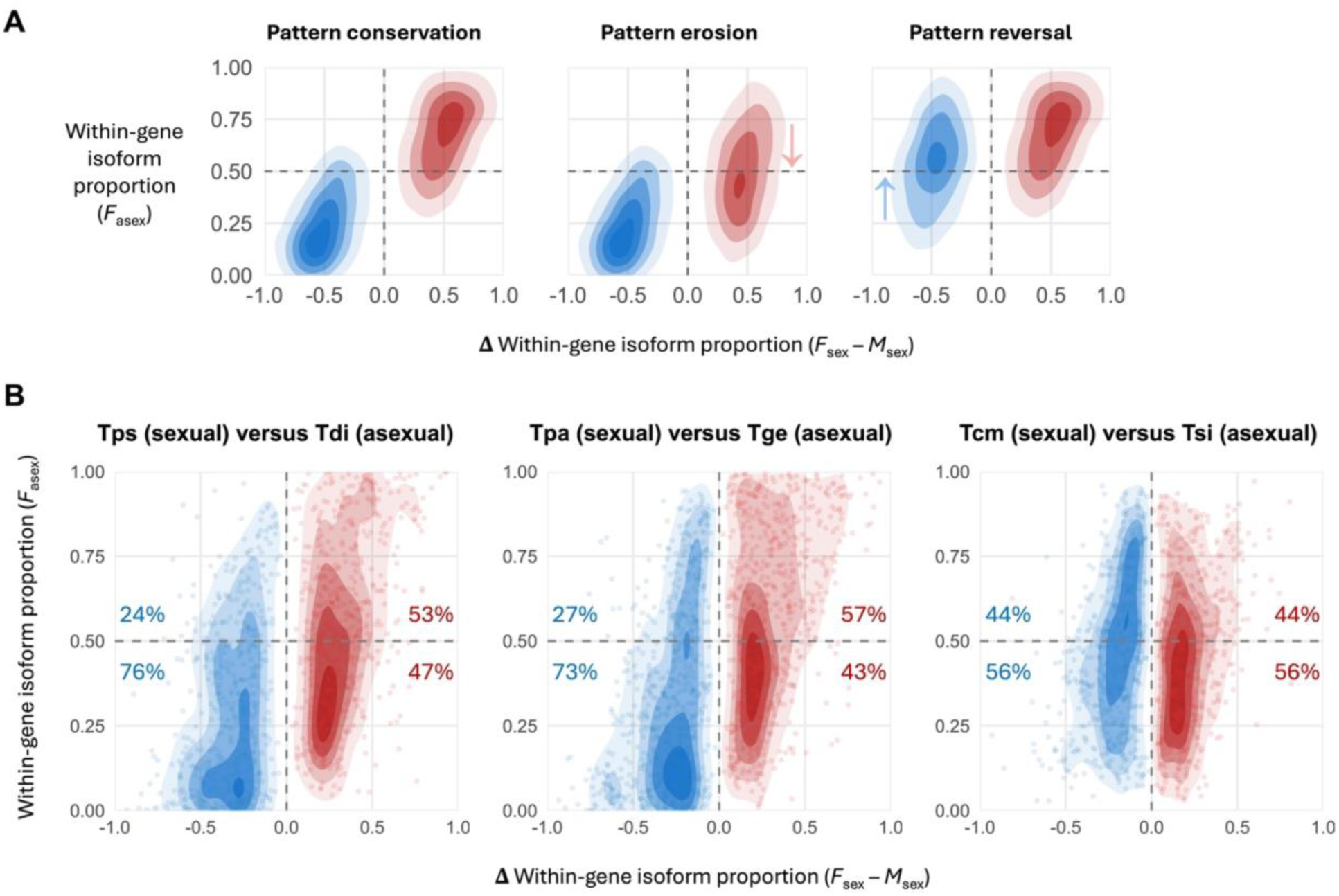
Changes in gonadal within-gene isoform proportions following the transition to asexuality. (A) Potential scenarios for patterns of relative isoform proportions in asexual females for isoforms identified as female-biased (red) and male-biased (blue) in sexual lineages. Under pattern conservation, female-biased isoforms maintain a relatively high usage (relative proportion) in asexual females, whereas male-biased isoforms remain at low proportions. Under pattern erosion, female-biased isoforms decrease in relative usage in asexual females, weakening the association between sexual sex bias and asexual female isoform proportion. Under pattern reversal, male-biased isoforms increase in relative proportions in asexual females, resulting in a reversal of the sexual female associated patterns. (B) Observed within-gene isoform proportions in asexual females relative to isoform sex bias in sexual species, across three sexual-asexual lineage comparisons. Each point represents an isoform, with female-biased isoforms in red and male-biased isoforms in blue. Shaded contours show two-dimensional kernel-density estimates, with darker shadings indicating greater density of isoforms. Percentages indicate the proportion of female-biased (red) and male-biased (blue) isoforms above and below 50% within-gene proportion in asexual females. Species name abbreviations: Tps = *T. poppense*, Tdi = *T. douglasi*, Tpa = *T. podura*, Tge = *T. genevievae*, Tcm = *T. californicum*, Tsi = *T. shepardi*.

Empirical distributions across three sexual-asexual lineage pairs predominantly support a combination of conservation and pattern erosion (Fig. 5B). The majority of ancestral female-and male-biased isoforms maintain high and, respectively, low within-gene proportions in asexual females, in line with the notion that regulatory conservation or broad developmental constraints preserve ancestral splicing patterns even after the transition to asexuality. Alternatively, limited time since the divergence between sexual and asexual lineages may have constrained the accumulation of changes in alternative splicing patterns.

Asexual females generally do not exhibit widespread reversal or masculinization of alternative splicing, with more than 70% of male-biased isoforms retaining low within-gene usage (exception being *T. shepardi*, where only 56% of male-biased isoforms have low relative proportions). In contrast, female-biased isoforms consistently show evidence of pattern erosion, with an average of 49% shifting towards low relative use in asexual females. One-sided exact binomial sensitivity analyses showed that the observed conservation of ancestral within-gene proportions among female-biased isoforms was significantly lower than conservation expectations of 55% or greater (*p* < 0.05, Table S3). Moreover, the probability of retaining the ancestral sex-associated isoform proportions significantly differed between female-and male-biased isoforms (one-sided Fisher’s exact tests, *p* < 0.001, Table S4), supporting preferential erosion of female-biased isoform usage following the transition to asexuality. We observe a similar splicing pattern erosion in the case of asexual males, with on average 43% of male-biased isoforms shifting towards low relative use in asexual males (one-sided exact binomial tests, minimum rejected null probability 61%, *p* < 0.05, Table S5). While the proportions of male-biased isoforms decreased, this was not accompanied by a reciprocal increase in female-biased isoforms in asexual males (one-sided Fisher’s exact tests, *p* < 0.001, Table S6).

Taken together, our results show that asexuality is associated with a partial erosion of sex-biased alternative splicing patterns. In both asexual females and males, we observed a parallel attenuation of isoforms ancestrally associated with an individual’s phenotypic sex, without a compensatory gain in opposite-sex isoforms. This parallel reduction in sex-concordant isoform usage suggests a generalized weakening of sex-specific splicing regulation following the transition to asexuality. Reduced sexual selection and sexual conflict, together with reduced efficacy of purifying selection and increased drift in asexual lineages, could have contributed to this pattern. Nevertheless, the comparable erosion of female-and male-associated splicing patterns does not support a simple expectation that relaxed selection on male function is the primary underlying factor, since this model predicts a particularly pronounced decay of male-biased isoforms in asexual males. In addition, we found no evidence in asexuals of increased splicing noise among high-confidence isoforms, arguing against a pervasive stochastic breakdown of splicing regulation. These findings suggest that sexual selection and divergent sex-specific phenotypic optima may actively maintain the complexity and regulatory differentiation of alternative splicing programs in sexual *Timema*, whereas their relaxation following the transition to asexuality permits partial erosion of sex-concordant isoform usage.

## Concluding Remarks

Alternative splicing is a pervasive and structured component of transcriptomic regulation in stick insects, with splicing patterns that vary across tissues, sexes, and evolutionary lineages.

By leveraging long-read transcriptomics across multiple species, tissues, sexes, and reproductive modes, we provide a comprehensive isoform-resolved view of splicing regulation in this system. Our results reveal several key insights. First, stick insects exhibit high levels of isoform diversity, particularly in gonadal tissue, matching or exceeding those reported in other invertebrates. We found no evidence for a large, genome-wide increase in isoform variation in asexuals, although our limited phylogenetically structured comparisons cannot rule out subtler effects of relaxed selection or drift. Instead, differences in isoform diversity were concentrated in gonadal tissue and sex-associated splicing contexts, a pattern consistent with alternative splicing contributing to reproductive and sex-specific transcriptomic regulation. Second, when considering the number of testable genes, sex-specific alternative splicing is as prevalent as sex-biased gene expression in gonads, indicating that splicing is not a secondary or minor contributor to sexual dimorphism, but a core regulatory mechanism. Despite rapid lineage-specific turnover, we identify a conserved core of differentially spliced genes, pointing to deep evolutionary constraints on key components of sexual differentiation. Third, differential splicing and differential expression act on largely distinct gene sets and are subject to different selective regimes, whereby splicing preferentially targets more broadly expressed, slowly evolving genes under strong purifying selection, consistent with a role in resolving sexual conflict without altering overall gene dosage. Finally, by comparing sexual and asexual lineages, we show that some sex-specific splicing patterns are eroded following the loss of sexual reproduction. Contrary to expectations under a simple drift-based model, asexual *Timema* do not exhibit increased splicing noise, but instead show reduced isoform diversity and a systematic loss of some of the sex-specialized isoforms. This pattern suggests that sexual selection and sex-specific optima may actively maintain splicing complexity in sexual species. Taken together, our study establishes alternative splicing as a central, evolutionarily dynamic, yet a selectively constrained layer of gene regulation that exhibits extensive sexual dimorphism and evolves in association with reproductive mode. It also highlights the power of long-read transcriptomics to uncover regulatory variation, and positions stick insects as a valuable model for dissecting the evolutionary forces governing transcriptome complexity.

## Acknowledgements

This work was supported by the European Molecular Biology Organization (EMBO) (grant number 167-2022 to I.D.). We thank the Lausanne Genomic Technologies Facilities for PacBio Iso-seq library preparation and sequencing. We would also like to thank members of the Schwander group and Linley M. Sherin for helpful comments and discussions on this work.

## Author Contributions

I.D. and T.S. designed research; I.D., M.L., V.M., and D.J.P. performed research; I.D. analyzed data with input from T.S.; I.D. and T.S. wrote the paper with input from all authors.

## Data availability

Iso-seq reads are available under NCBI BioProject PRJNA1149183. Scripts for data processing and analysis are available at https://github.com/idarolti/Darolti_etal_2026_AltSpli.

