## Supplementary Materials for "Sexually dimorphic alternative splicing and its erosion following the loss of sex in stick insects"

**Table S1. Sample information for Iso-seq library preparation.**

| Species Pairs | Species | Reproductive Mode | Sex | Tissue |
|---|---|---|---|---|
| Pair 1 | *T. poppense* (Tps) | Sexual | Female | Femur<br>Gut<br>Gonad |
| | | | Male | Femur<br>Gut<br>Gonad |
| | *T. douglasi* (Tdi) | Asexual | Female | Femur<br>Gut<br>Gonad |
| | | | Male | Femur<br>Gut<br>Gonad |
| Pair 2 | *T. californicum* (Tcm) | Sexual | Female | Femur<br>Gut<br>Gonad |
| | | | Male | Femur<br>Gut<br>Gonad |
| | *T. shepardi* (Tsi) | Asexual | Female | Femur<br>Gut<br>Gonad |
| Pair 3 | *T. cristinae* (Tce) | Sexual | Female | Femur<br>Gut<br>Gonad |
| | | | Male | Femur<br>Gut<br>Gonad |
| | *T. monikensis* (Tms) | Asexual | Female | Femur<br>Gut |
| | | | Male | Femur<br>Gut<br>Gonad |
| Pair 4 | *T. podura* (Tpa) | Sexual | Female | Femur<br>Gut<br>Gonad |
| | | | Male | Femur<br>Gut<br>Gonad |
| | *T. genevievae* (Tge) | Asexual | Female | Femur<br>Gut<br>Gonad |
| | | | Male | Femur<br>Gut<br>Gonad |
| - | *T. bartmani* (Tbi) | Sexual | Female | Gut<br>Gonad |
| | | | Male | Gut<br>Gonad |

**Table S2. Gene counts, isoform counts, and splice type events for each assembled transcriptome.**

| Transcriptome | Number of genes | Number of filtered isoforms | Splicing events | | | | |
|---|---|---|---|---|---|---|---|
| | | | Exon Skipping | Intron Retention | Alternative 5’ or 3’ end | Novel Splice Site | Other* |
| *T. poppense* + *T. douglasi* | 12,681 | 30,212 | 2,060 | 1,060 | 2,916 | 11,875 | 4,962 |
| *T. californicum* + *T. shepardi* | 11,736 | 22,225 | 869 | 596 | 3,107 | 6,651 | 1,633 |
| *T. cristinae* + *T. monikensis* | 11,863 | 22,131 | 805 | 578 | 2,585 | 6,200 | 2,234 |
| *T. podura* + *T. genevievae* | 12,229 | 25,926 | 953 | 849 | 2,777 | 9,364 | 2,966 |
| *T. bartmani* | 10,891 | 18,686 | 1,418 | 785 | 2,618 | 3,363 | 850 |

*as defined by the SQANTI3 classification, this category includes novel genes, fused genes, antisense isoforms (do not overlap a same-strand reference gene), genic isoforms (isoforms completely contained in an intronic sequence or partially overlapping an intron or exon)

**Table S3. Sensitivity analysis of null expectations for conservation of sex-biased isoform proportions in asexual females.** Species name abbreviations: Tps = *T. poppense*, Tdi = *T. douglasi*, Tcm = *T. californicum*, Tsi = *T. shepardi*, Tpa = *T. podura*, Tge = *T. genevievae*.

| **Lineage pair** | **Female-biased isoforms** | **High asexual female-biased isoform proportion (>0.5)** | **Percentage female-biased isoforms with conserved ancestral pattern** | **Minimum rejected null probability (exact Binomial test, *p* <0.05)** |
|---|---|---|---|---|
| Tps – Tdi | 529 | 281 | 53% | 57% |
| Tpa – Tge | 704 | 398 | 57% | 60% |
| Tcm – Tsi | 554 | 246 | 44% | 48% |

**Table S4. Fisher's exact test results for differences between the conservation of female-biased and male-biased isoforms in asexual females.** Species name abbreviations: Tps = *T. poppense*, Tdi = *T. douglasi*, Tcm = *T. californicum*, Tsi = *T. shepardi*, Tpa = *T. podura*, Tge = *T. genevievae*.

| **Lineage pair** | **Odds ratio for conservation, female- vs. male-biased** | **One-sided 95% CI** | **One-sided Fisher's exact *p* value** |
|---|---|---|---|
| Tps – Tdi | 0.35 | 0.00 – 0.44 | < 0.001 |
| Tpa – Tge | 0.47 | 0.00 – 0.58 | < 0.001 |
| Tcm – Tsi | 0.63 | 0.00 – 0.78 | < 0.001 |

**Table S5. Sensitivity analysis of null expectations for conservation of sex-biased isoform proportions in asexual males.** Species name abbreviations: Tps = *T. poppense*, Tdi = *T. douglasi*, Tce = *T. cristinae*, Tms = *T. monikensis*, Tpa = *T. podura*, Tge = *T. genevievae*.

| **Lineage pair** | **Male-biased isoforms** | **High asexual male-biased isoform proportion (>0.5)** | **Percentage male-biased isoforms with conserved ancestral pattern** | **Minimum rejected null probability (exact Binomial test, *p* <0.05)** |
|---|---|---|---|---|
| Tps – Tdi | 499 | 299 | 60% | 64% |
| Tpa – Tge | 692 | 388 | 56% | 59% |
| Tce – Tms | 578 | 329 | 57% | 60% |

**Table S6. Fisher's exact test results for differences between the conservation of female-biased and male-biased isoforms in asexual males.** Species name abbreviations: Tps = *T. poppense*, Tdi = *T. douglasi*, Tce = *T. cristinae*, Tms = *T. monikensis*, Tpa = *T. podura*, Tge = *T. genevievae*.

| **Lineage pair** | **Odds ratio for conservation, female- vs. male-biased** | **One-sided 95% CI** | **One-sided Fisher's exact *p* value** |
|---|---|---|---|
| Tps – Tdi | 0.35 | 0.00 – 0.45 | < 0.001 |
| Tpa – Tge | 0.47 | 0.00 – 0.57 | < 0.001 |
| Tce – Tms | 0.60 | 0.00 – 0.75 | < 0.001 |

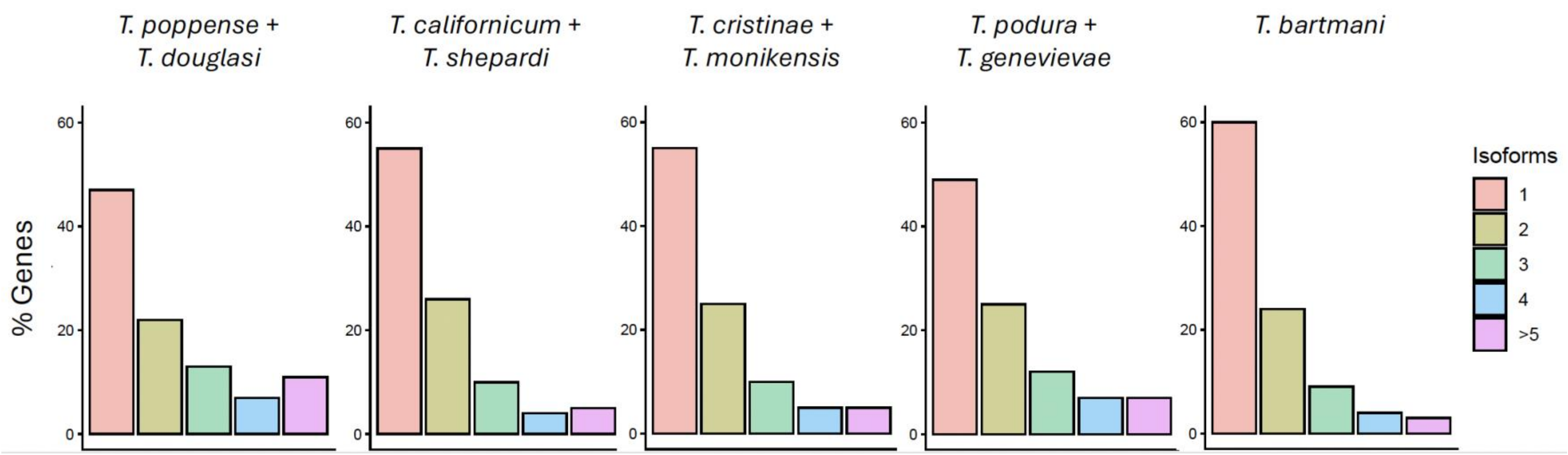


**Fig. S1. Per-gene isoform diversity for each *Timema* transcriptome assembly.** On average across species, 47% of genes have two or more expressed isoforms.

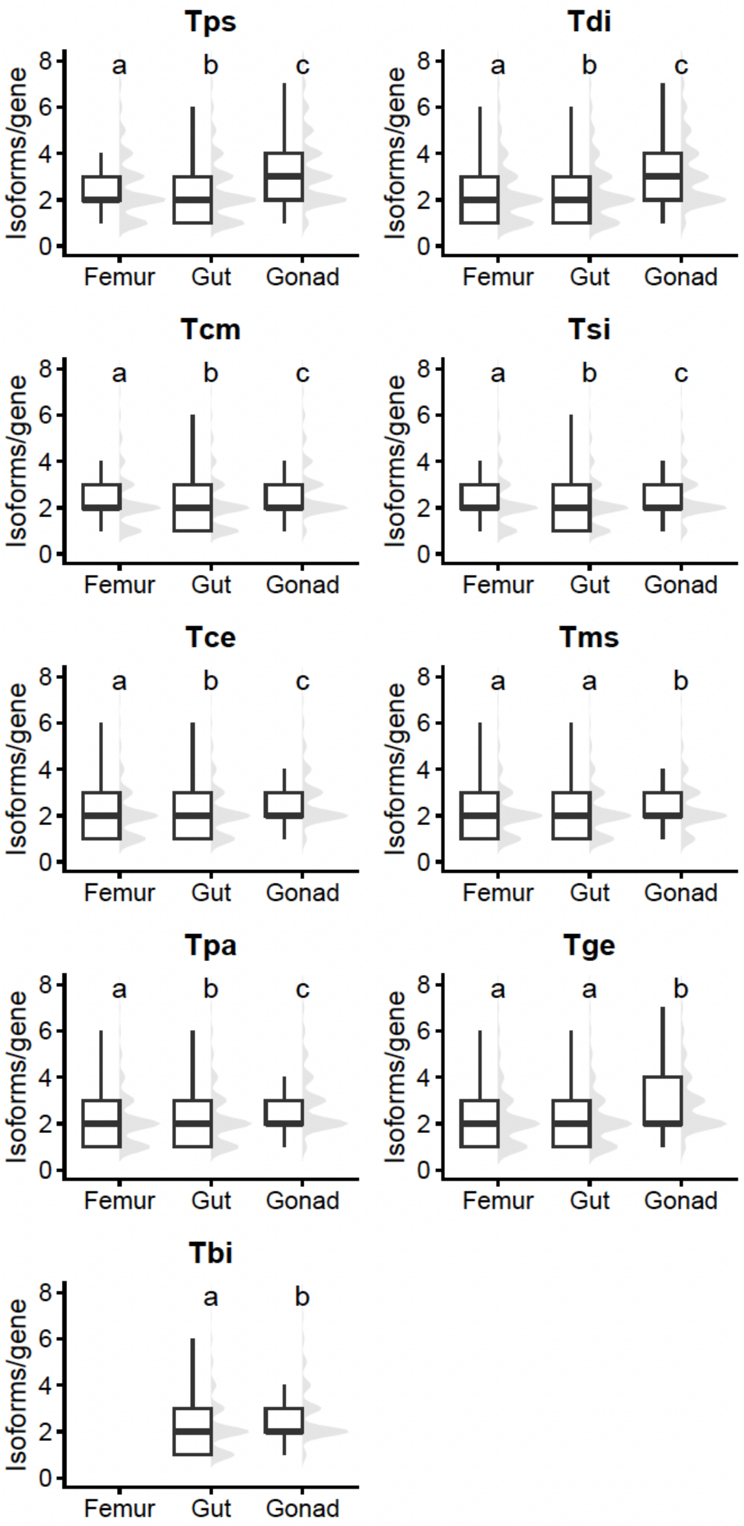


**Fig. S2. Distribution of number of isoforms per gene in each tissue of each *Timema* species.** For each species plot, only genes expressed in all tissues and those with more than one isoform in at least one of the tissues are included. Results from pairwise Wilcoxon signed-rank tests are indicated by the lowercase letters, where boxplots labelled with different letters are significantly different from each other ($p_{adj} < 0.05$), while boxplots sharing at least one letter are non-significant. Species name abbreviations: Tps = *T. poppense*, Tdi = *T. douglasi*, Tcm = *T. californicum*, Tsi = *T. shepardi*, Tce = *T. cristinae*, Tms = *T. monikensis*, Tpa = *T. podura*, Tge = *T. genevievae*, Tbi = *T. bartmani*.

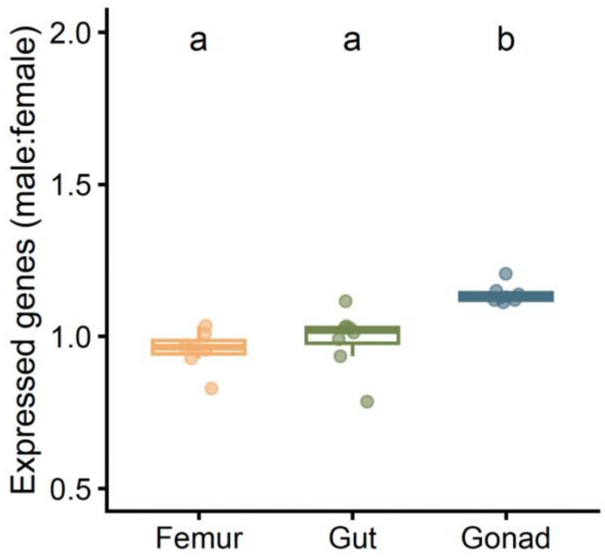


**Fig. S3. The ratio between males and females in the number of expressed genes across all *Timema* species.** Results from pairwise Wilcoxon rank-sum tests are represented by lowercase letters, where boxplots labelled with distinct letters are significantly different ($p_{adj}$ < 0.05), while boxplots sharing a letter are non-significant.

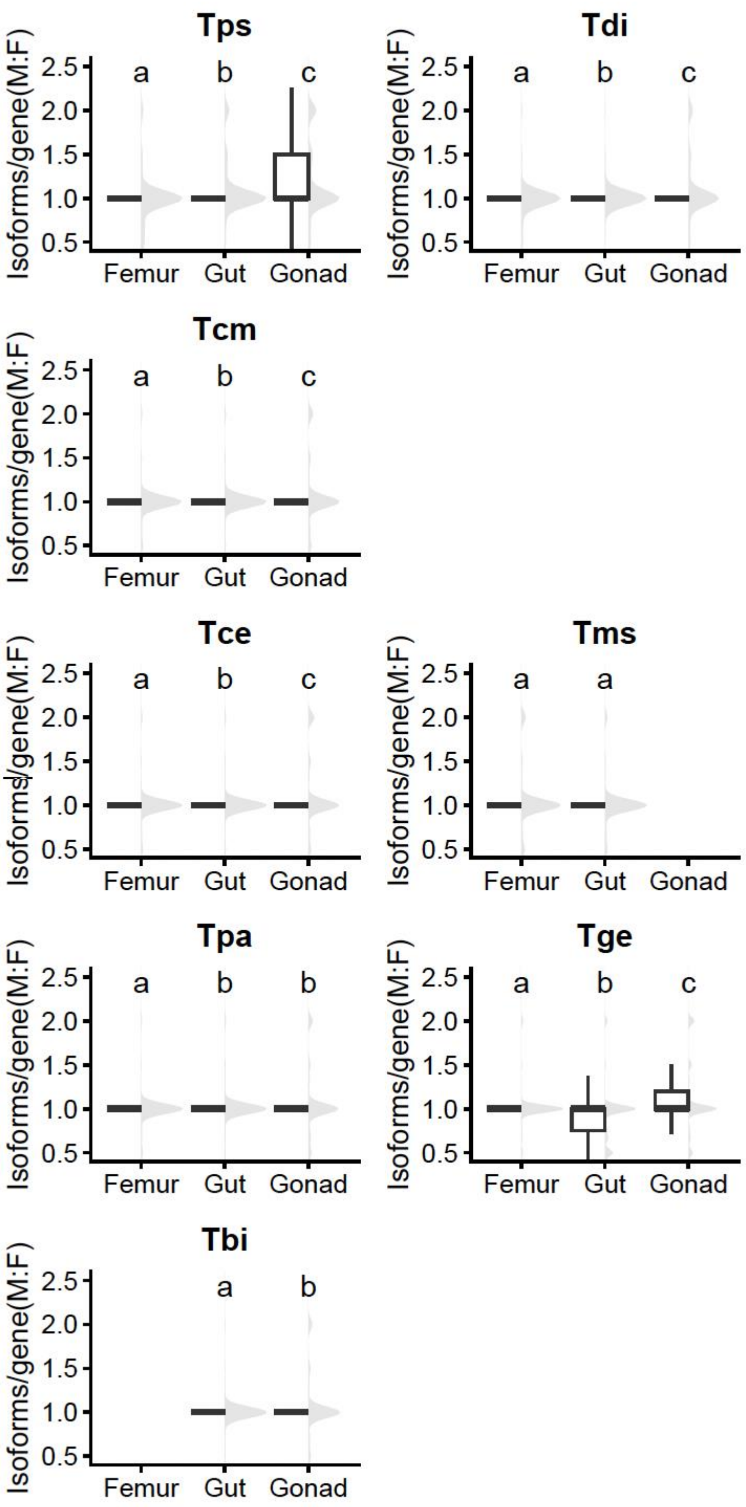


**Fig. S4. Distribution of the ratio between males and females in the number of isoforms per gene for each tissue and species.** For each species plot, only genes expressed in both males and females and those with at more than one isoform in at least one of the sexes are included. Results from pairwise Wilcoxon rank-sum tests are indicated by the lowercase letters, where boxplots labelled with different letters are significantly different from each other ($p_{adj} < 0.05$), while boxplots sharing at least one letter are non-significant. Species name abbreviations: Tps = *T. poppense*, Tdi = *T. douglasi*, Tcm = *T. californicum*, Tsi = *T. shepardi*, Tce = *T. cristinae*, Tms = *T. monikensis*, Tpa = *T. podura*, Tge = *T. genevievae*, Tbi = *T. bartmani.*

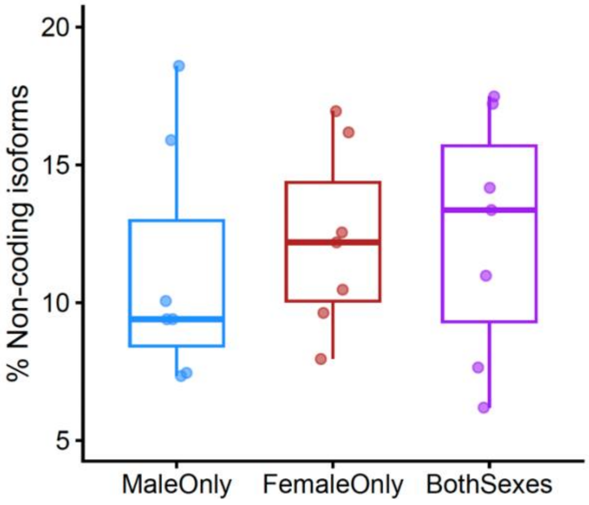


**Fig. S5. Percentage of isoforms that are non-coding out of the total male-specific isoforms (blue), female-specific isoforms (red), and isoforms expressed in both sexes (purple).** Data refers to the gonadal tissue of all species. Wilcoxon rank-sum tests indicate no significant differences in either of the pairwise comparisons ($p_{adj} > 0.05$).

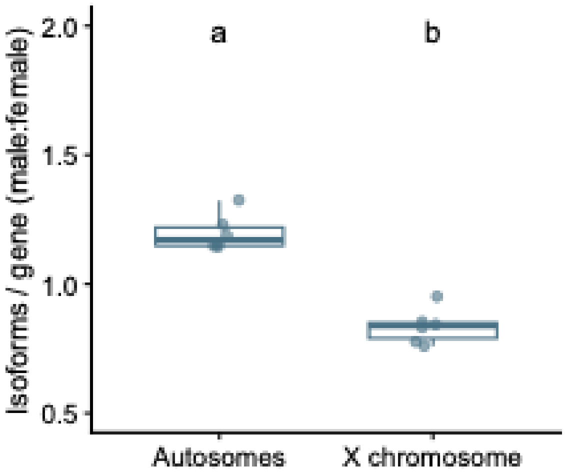


**Fig. S6. Ratio of per-gene isoform diversity between males and females for autosomal and X-lined gonadal expressed genes.** Only genes expressed in both sexes are included, and genes with only one isoform in both males and females have been excluded. Significant difference ($p_{adj} < 0.05$) based on Wilcoxon rank-sum tests between autosomal and X-linked genes is represented by the lowercase letters.

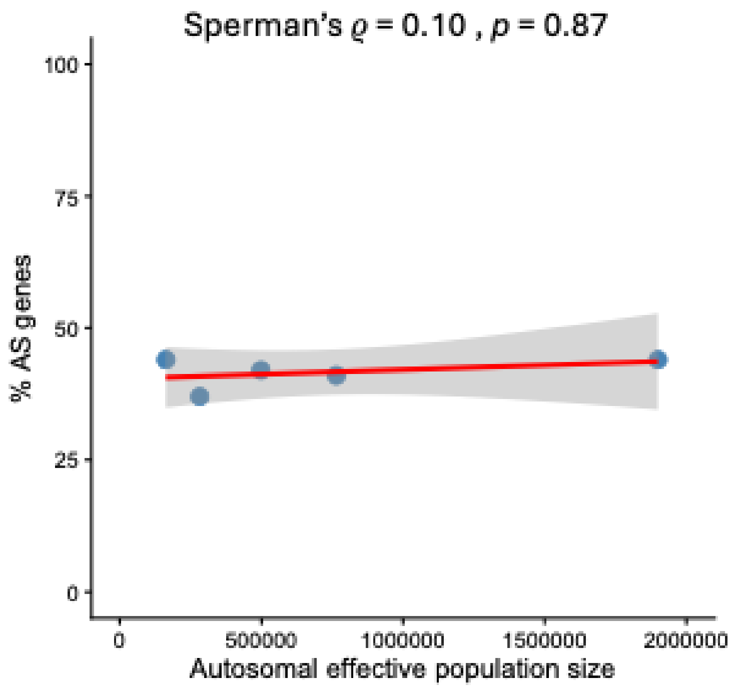


**Fig. S7. Correlation between the autosomal effective population size (*Ne*) of each sexual *Timema* species and the percentage of alternatively spliced genes.** Estimates of *Ne* are derived from Parker et al. 2022. The isoform analysis is based on a randomized subsampled dataset of 400,000 FLNCs from each library, and on genes expressed in all species.

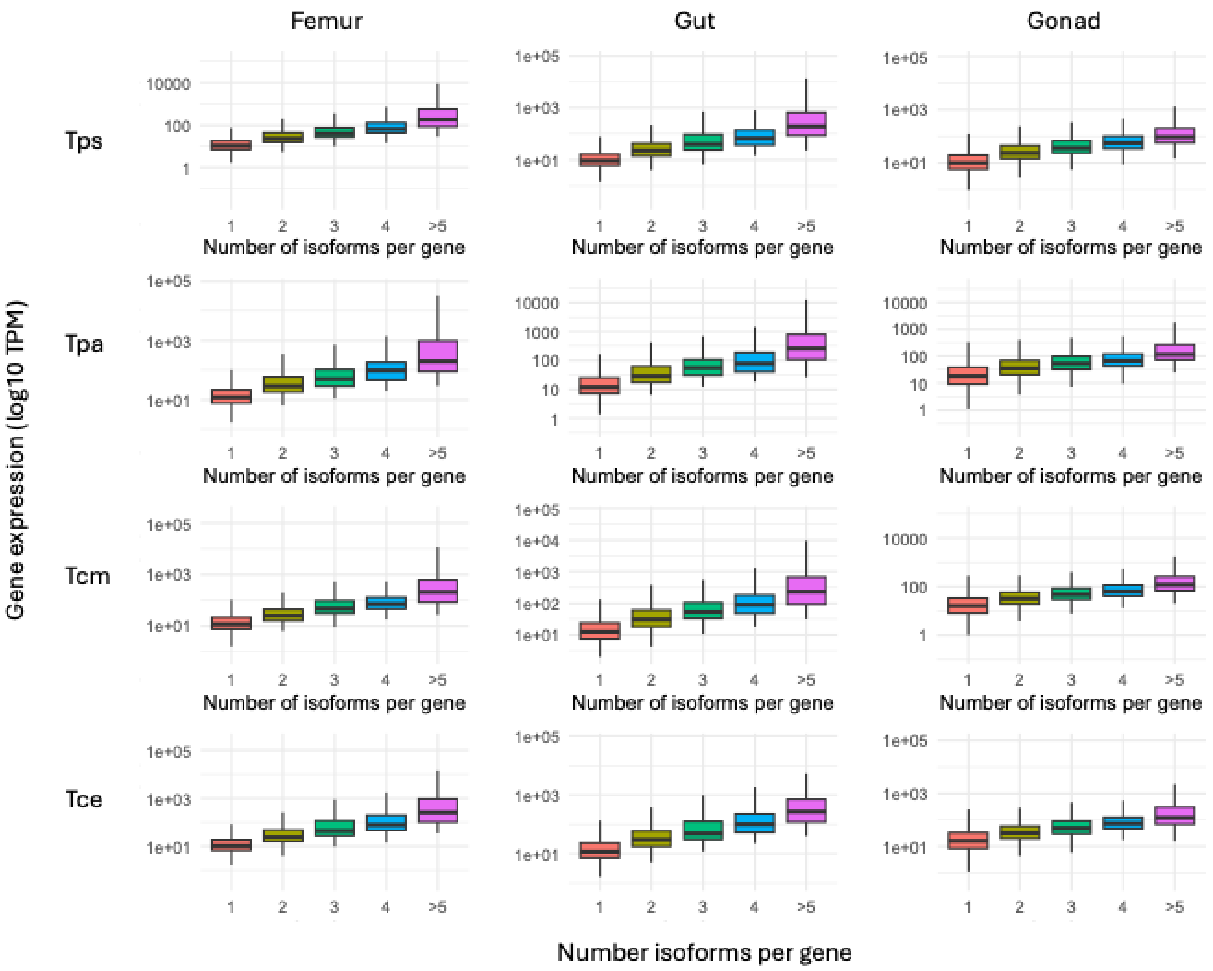


**Fig. S8. Gene expression estimates for genes grouped by their number of isoforms.** Species name abbreviations: Tps = *T. poppense*, Tcm = *T. californicum*, Tce = *T. cristinae*, Tpa = *T. podura.*

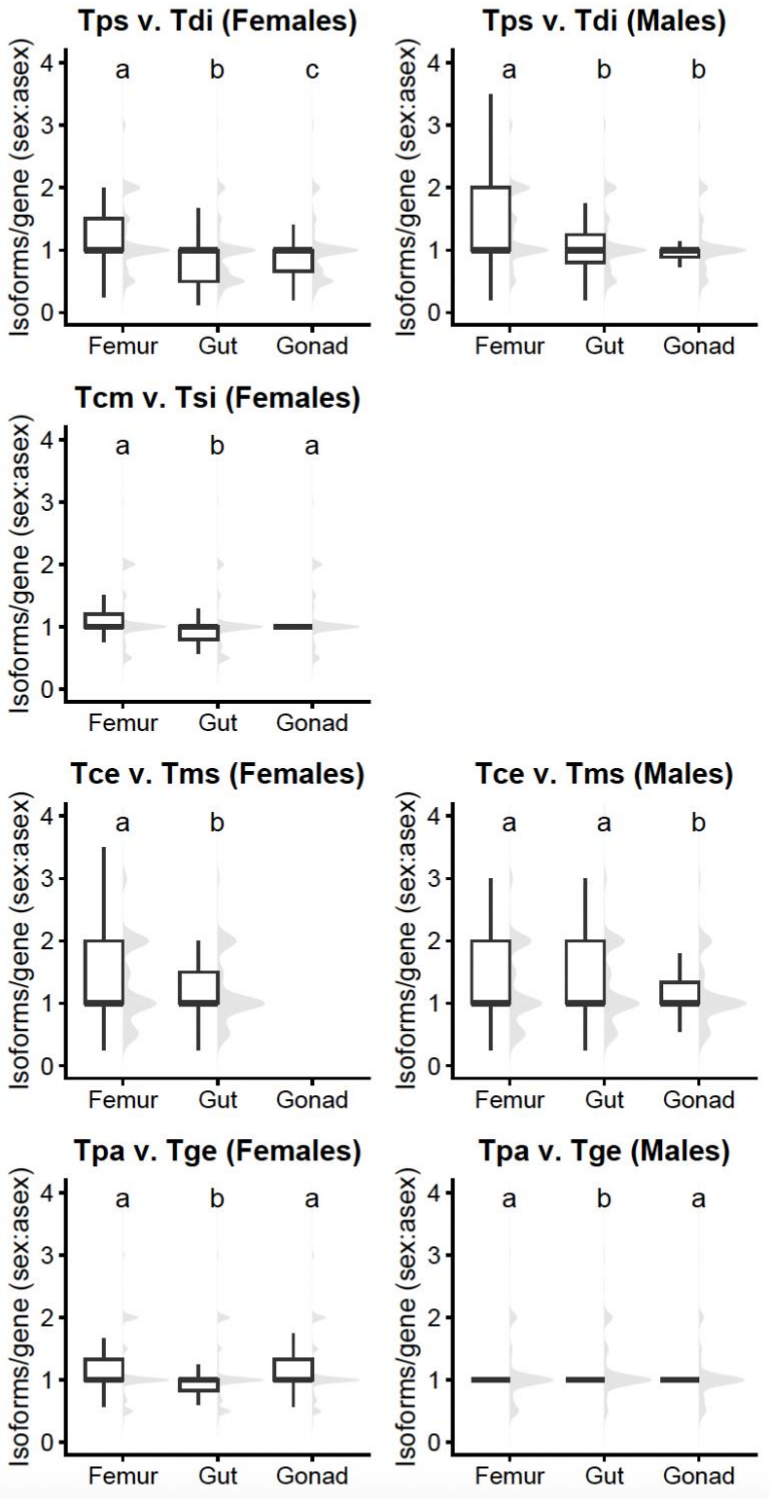


**Fig. S9. Distribution of the ratio between sexual and asexual individuals in the number of isoforms per gene for each tissue.** Only genes expressed in both sexual and asexual individuals and those that have more than one isoform in at least one reproductive mode are included. Results from pairwise Wilcoxon rank-sum tests are indicated by the lowercase letters, where boxplots labelled with different letters are significantly different from each other ($p_{adj} < 0.05$), while boxplots sharing at least one letter are non-significant. Species name abbreviations: Tps = *T. poppense*, Tdi = *T. douglasi*, Tcm = *T. californicum*, Tsi = *T. shepardi*, Tce = *T. cristinae*, Tms = *T. monikensis*, Tpa = *T. podura*, Tge = *T. genevievae*.

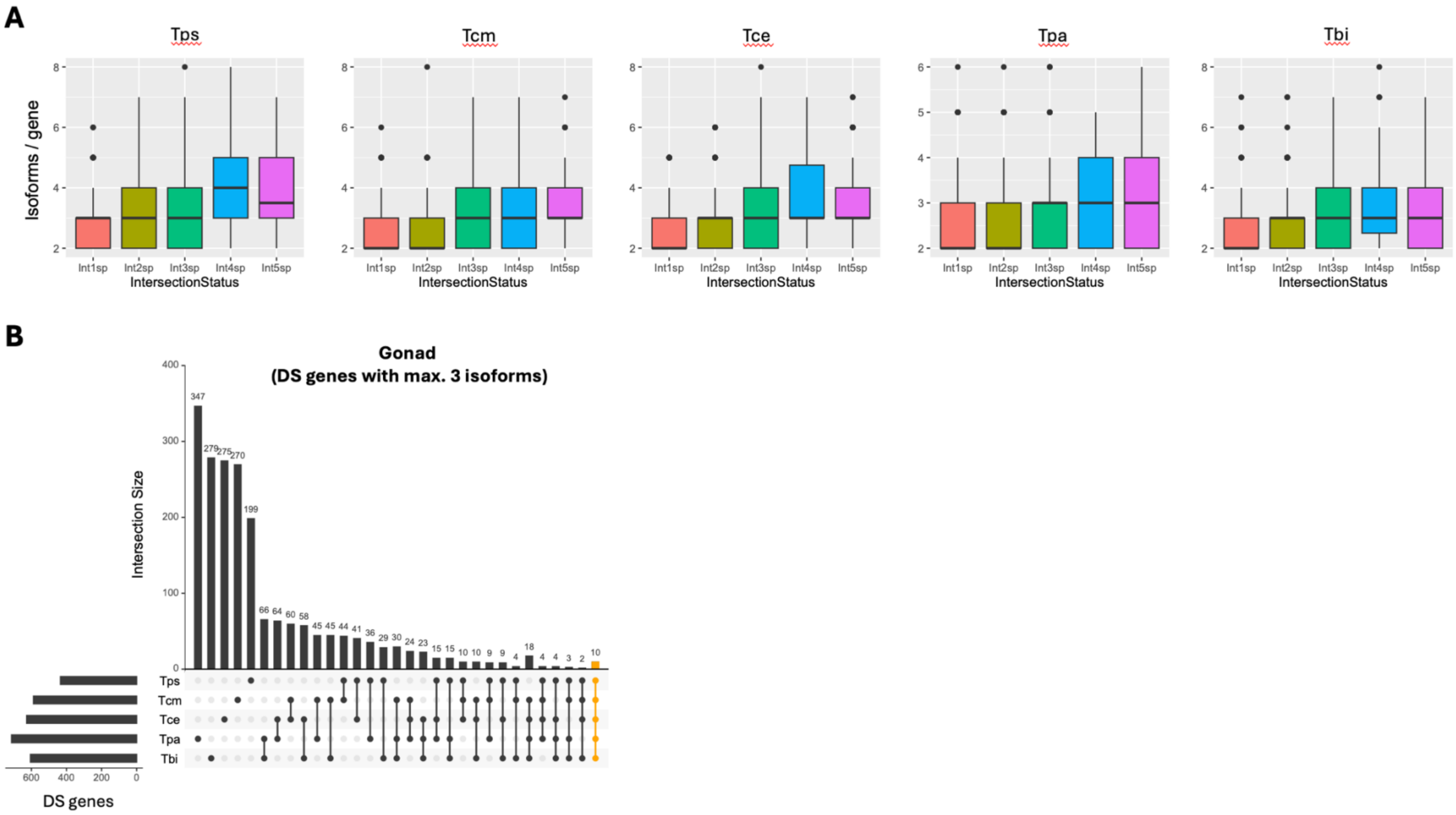


**Fig. S10**. (A) Relationship between gene complexity, measured as the number of isoforms per gene, and the conservation of differential splicing (DS) patterns, ranging from species-specific DS genes (Int1sp) to genes with DS status conserved across all five species (Int5sp). (B) Intersection of DS genes across the five sexual *Timema* species, for orthologous gonad-expressed genes with a maximum of three isoforms. Species name abbreviations: Tps = *T. poppense*, Tcm = *T. californicum*, Tce = *T. cristinae*, Tpa = *T. podura*, Tbi = *T. bartmani*

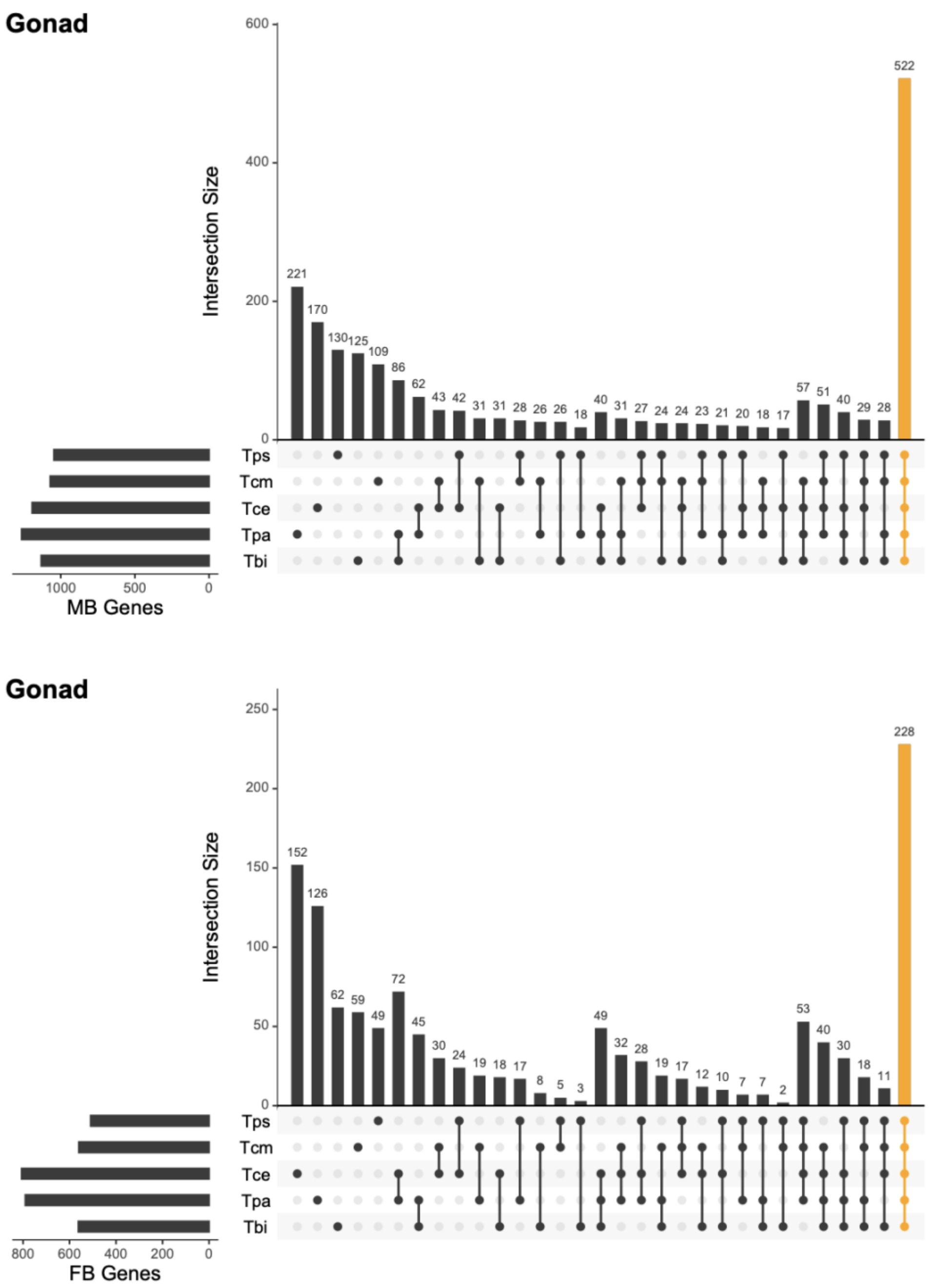


**Fig. S11.** Intersection of orthologous gonad male-biased (MB) and female-biased (FB) genes across the five sexual *Timema* species. Species name abbreviations: Tps = *T. poppense*, Tcm = *T. californicum*, Tce = *T. cristinae*, Tpa = *T. podura*, Tbi = *T. bartmani*

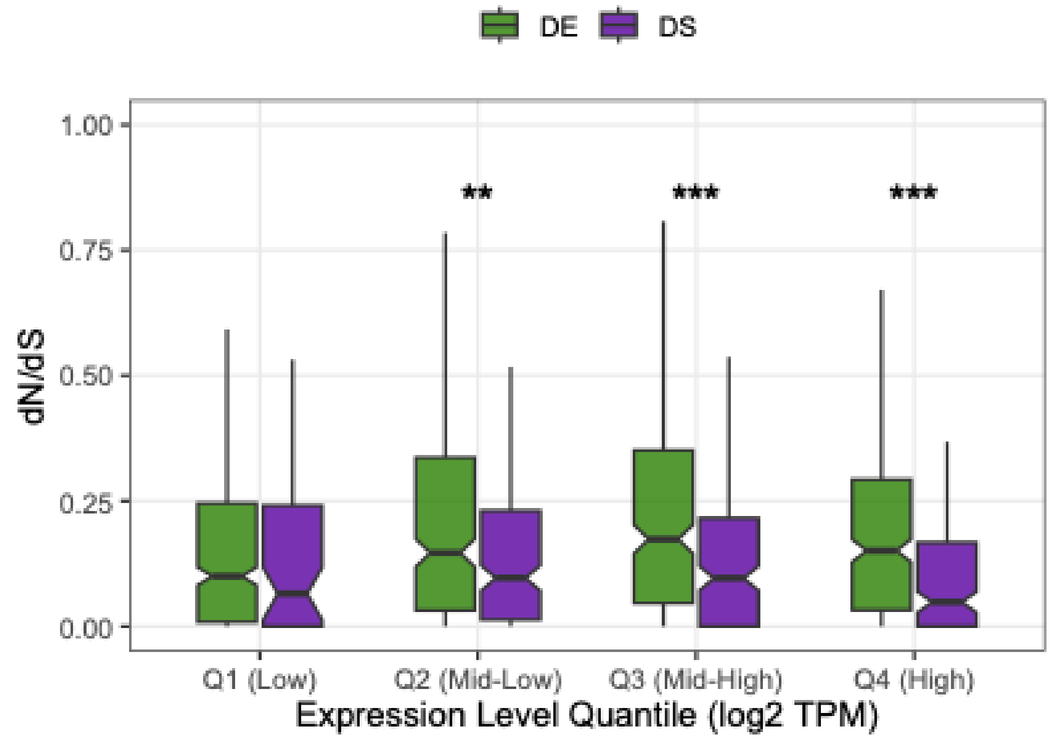


**Fig. S12. Evolutionary rate ($d_N/d_S$) of differentially expressed (DE) and differentially spliced (DS) genes across expression quartiles.** Statistical significance between DE and DS genes within each quartile was assessed using Wilcoxon rank-sum tests with BH FDR adjustment (** $p_{adj} < 0.01$, *** $p_{adj} < 0.001$). Sample sizes ($n_{DE}$, $n_{DS}$): Q1 (428, 53), Q2 (292, 189), Q3 (283, 1980), Q4 (319, 162).

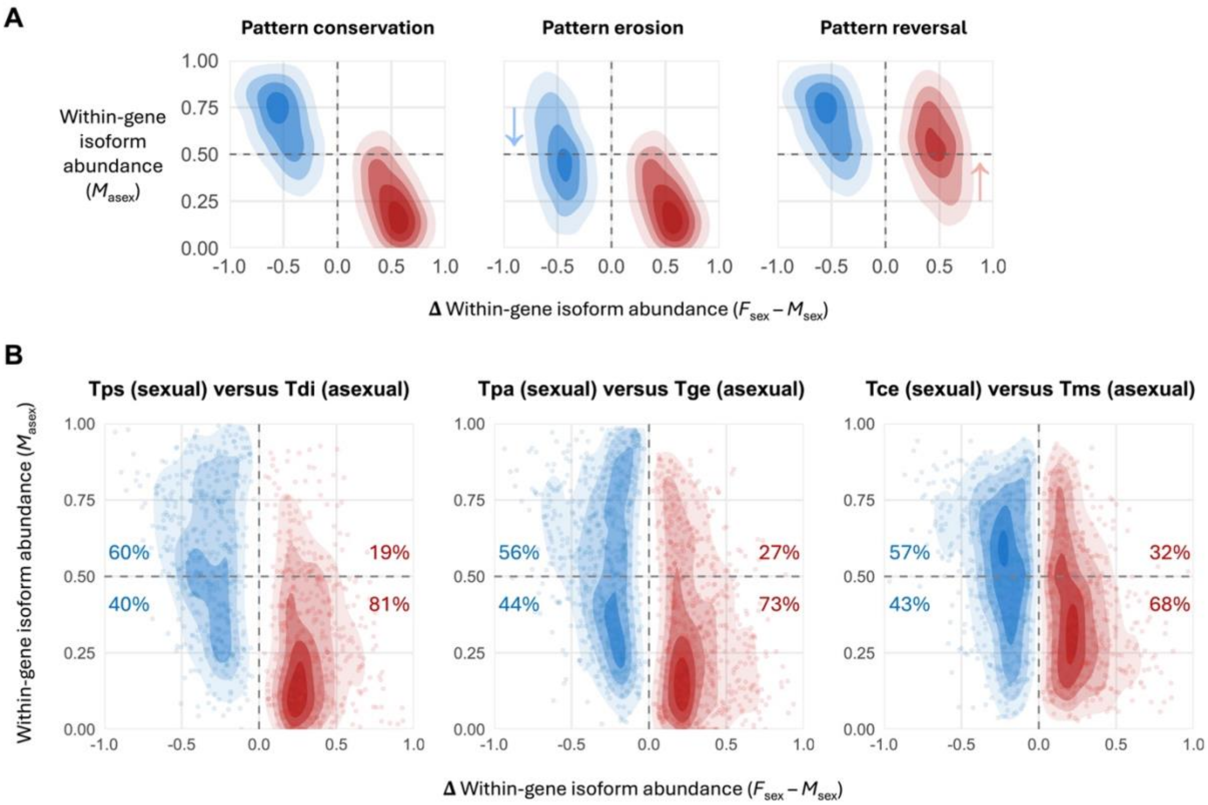


**Fig. S13. Changes in gonadal within-gene isoform proportions following the transition to asexuality.** (A) Potential scenarios of patterns of relative isoform proportions in asexual males for isoforms identified as female-biased (red) and male-biased (blue) in sexual lineages. Under pattern conservation, male-biased isoforms maintain a relatively high usage (relative proportion) in asexual males, whereas female-biased isoforms remain at low proportions. Under pattern erosion, male-biased isoforms decrease in relative usage in asexual males, weakening the association between sexual sex bias and asexual male isoform proportion. Under pattern reversal, female-biased isoforms increase in relative proportions in asexual males, resulting in a reversal of the sexual male associated patterns. (B) Observed within-gene isoform proportions in asexual males relative to isoform sex bias in sexual species, across three sexual-asexual lineage comparisons. Each point represents an isoform, with female-biased isoforms in red and male-biased isoforms in blue. Shaded contours show two-dimensional kernel-density estimates, with darker shadings indicating greater density of isoforms. Percentages indicate the proportion of female-biased (red) and male-biased (blue) isoforms above and below 50% within-gene proportion in asexual males. Species name abbreviations: Tps = *T. poppense*, Tdi = *T. douglasi*, Tpa = *T. podura*, Tge = *T. genevievae*, Tce = *T. cristinae*, Tms = *T. monikensis*.
